# The Amphibian Antimicrobial Peptide Uperin 3.5 is a Cross-α/Cross-β Chameleon Functional Amyloid

**DOI:** 10.1101/2020.05.31.126045

**Authors:** Nir Salinas, Einav Tayeb-Fligelman, Massimo Sammito, Daniel Bloch, Raz Jelinek, Dror Noy, Isabel Uson, Meytal Landau

**Author notes:** These authors contributed equally (Nir Salinas, Einav Tayeb-Fligelman).

## Abstract

Antimicrobial activity is being increasingly linked to amyloid fibril formation, suggesting physiological roles for some human amyloids, which have historically been viewed as strictly pathological agents. This work reports on formation of functional cross-α amyloid fibrils of the amphibian antimicrobial peptide uperin 3.5 at atomic-resolution, an architecture initially discovered in the bacterial PSMα3 cytotoxin. The fibrils of uperin 3.5 and PSMα3 were comprised of parallel and anti-parallel helical sheets, respectively, recapitulating properties of β-sheets. Uperin 3.5 helical fibril formation was largely induced by, and formed on, bacterial cells or membrane mimetics, and led to membrane damage and cell death. Uperin 3.5 demonstrated chameleon properties, with a secondary structure switch to cross-β fibrils with reduced antibacterial activity in the absence of lipids or after heat shock. These findings suggest a regulation mechanism, which includes storage of inactive peptides as well as environmentally induced activation of uperin 3.5, via chameleon cross-α/β amyloid fibrils.

## Introduction

Antimicrobial peptides (AMPs) are found in all kingdoms of life, serving roles in the host-defense system by fighting microbial infections and killing cancerous cells^1–4^. AMPs mainly target and disrupt membranes, leading to cell death^1^, and, in some cases, their self-assembly into supramolecular structures enhances antimicrobial activity^5^. Specifically, certain AMPs, such as dermaseptin S9, assemble into well-ordered fibrils that resemble amyloids, proteins which form a cross-β architecture comprised of tightly mated β-sheets^6–17^. Human amyloids have primarily been associated with neurodegenerative and systemic diseases^18,19^, and evidence of antimicrobial properties for some, including the Alzheimer’s associated amyloid-β and Parkinson’s associated α-synuclein, suggest a physiological role in fighting infections threatening the brain^6,20–26^.

Our earlier investigations of functional amyloids revealed a distinct structure-function correlation in the phenol-soluble modulins (PSMs) family of peptides secreted by the *Staphylococcus aureus* bacterium, which are involved in virulence activities^27,28^, and form amyloid fibrils with specific morphologies^29–31^. Specifically, the biofilm-associated PSMα1 and PSMα4 adopt the amyloid ultra-stable cross-β architecture^29^, likely to serve as a scaffold rendering the biofilm a more resistant barrier. Exceptionally, PSMα3, which plays roles in cytotoxicity against human immune cells^27,32^, forms cross-α amyloid fibrils that are composed entirely of amphipathic α-helices. The helices stack perpendicular to the fibril axis into mated ‘sheets’^31^, just as the β-strands assemble in amyloid cross-β fibrils. Furthermore, a short segment from PSMα3 forms atypical β-rich fibrils with antiparallel orientation and shows a mild antibacterial activity^29^. Overall, PSMαs show diverse activities as well as different morphologies of amyloid and amyloid-like fibrils. We previously showed that the ability of PSMα3 to form cross-α fibrils is critical for cytotoxicity, likely mediated through a dynamic process of co-aggregation with membrane lipids^31^. Recently, various synthetic peptides, unnatural enantiomers and protein-mimics have been shown to form an architecture resembling cross-α^33–37^.

Previous studies by Bowie, Carver, Martin and co-workers showed that the AMP uperin 3.5, secreted on the skin of *Uperoleia mjobergii* (Australian toadlet)^38^, forms amyloid fibrils, and suggested an interaction with bacterial membrane lipids that stabilize its α-helical conformation^39–42^. Moreover, by using uperin 3.5 mutants, they showed that a high α-helical content and lower net charge contribute to higher aggregation rate of the peptide^40,41^. Here, we demonstrate, at atomic resolution, that uperin 3.5 formed cross-α fibrils, which we suggest being essential for its toxic activity against the Gram-positive bacterium *Micrococcus luteus*. Moreover, the presence of bacterial membrane lipids induced a structural transition of uperin 3.5 into helical species in solution and in the fibrils, whereas their absence revealed a chameleon behavior of a secondary structure switch into cross-β fibrils, which correlated with reduced antibacterial activity.

## Results

### Uperin 3.5 is a functional cross-α amyloid

As was previously shown^39^, the antimicrobial peptide uperin 3.5 self-assembled to form elongated amyloid fibrils, as visualized using transmission electron-microscopy (TEM), which bound the amyloid indicator dye thioflavin-T (ThT) (Figure S1). The crystal structure of the full-length, 17-residue AMP, uperin 3.5, determined at 1.45 Å resolution (Table 1) (PDB code: 6GS3), revealed a cross-α fibril architecture (Figure 1). The cross-α fibrils of uperin 3.5 formed from stacks of amphipathic α-helices, positioned perpendicular to the fibril axis, creating a unique arrangement of mated ‘helical sheets’ extending along the fibril axis, similar to the β-sheets which form cross-β amyloid fibril architecture^6–17^. Overall, the cross-α fibrils reassemble cross-β fibrils in their quaternary structures and in the ability to induce ThT fluorescence.

**Table 1.** Data collection and refinement statistics.

| **Uperin 3.5 cross-α fibril** |  |
| --- | --- |
| PDB accession code | 6GS3 |
| Beamline | ESRF MASSIF-3 |
| Date | April 19, 2017 |
| **Data collection** |  |
| Space group | P 1 2<sub>1</sub> 1 |
| Cell dimensions |  |
| *a*, *b*, *c* (Å) | 19.70, 28.44, 20.32 |
| $\alpha$, $\beta$, $\gamma$ (°) | 90.00, 106.95, 90.0 |
| Wavelength (Å) | 0.968 |
| Resolution (Å) | 19.47-1.45 (1.49-1.45) |
| R-factor observed (%) | 8.5 (60.1) |
| <sup>a</sup> *R*<sub>meas</sub> (%) | 8.9 (64.8) |
| *I* / σ*I* | 15.7 (2.7) |
| Completeness (%) | 97.0 (94.7) |
| Redundancy | 10.6 (7.6) |
| <sup>b</sup> CC<sub>1/2</sub> (%) | 99.9 (90.8) |
| **Refinement** |  |
| Resolution (Å) | 19.44-1.45 (1.49-1.45) |
| Completeness (%) | 97.2 (94.7) |
| <sup>c</sup> No. reflections | 3398 (238) |
| <sup>d</sup> *R*<sub>work</sub> (%) | 18.5 (24.4) |
| *R*<sub>free</sub> (%) | 22.9 (28.0) |
| No. atoms | 275 |
| Protein | 254 |
| Ligand/ion | 5 |
| Water | 16 |
| *B*-factors |  |
| Protein | 15.1 (Chain A)<br>12.7 (Chain B) |
| Ligand/ion | 41.4 (K+)<br>19.3 (SCN-) |
| Water | 21.6 |
| R.m.s. deviations |  |
| Bond lengths (Å) | 0.016 |
| Bond angles (°) | 1.783 |
| Clash score<sup>80</sup> | 0 |
| Molprobity score<sup>80</sup> | 0.5 |
| Molprobity percentile<sup>80</sup> | 100th percentile |
| Number of crystals used for scaling | 1 |

Values in parentheses are for highest-resolution shell.
(<sup>a</sup>) *R*-meas is a redundancy-independent R-factor defined in<sup>81</sup>.
(<sup>b</sup>) CC<sub>1/2</sub> is percentage of correlation between intensities from random half-datasets<sup>82</sup>.
(<sup>c</sup>) Number of reflections corresponds to the working set.
(<sup>d</sup>) *R*work corresponds to working set.

**Figure 1.**
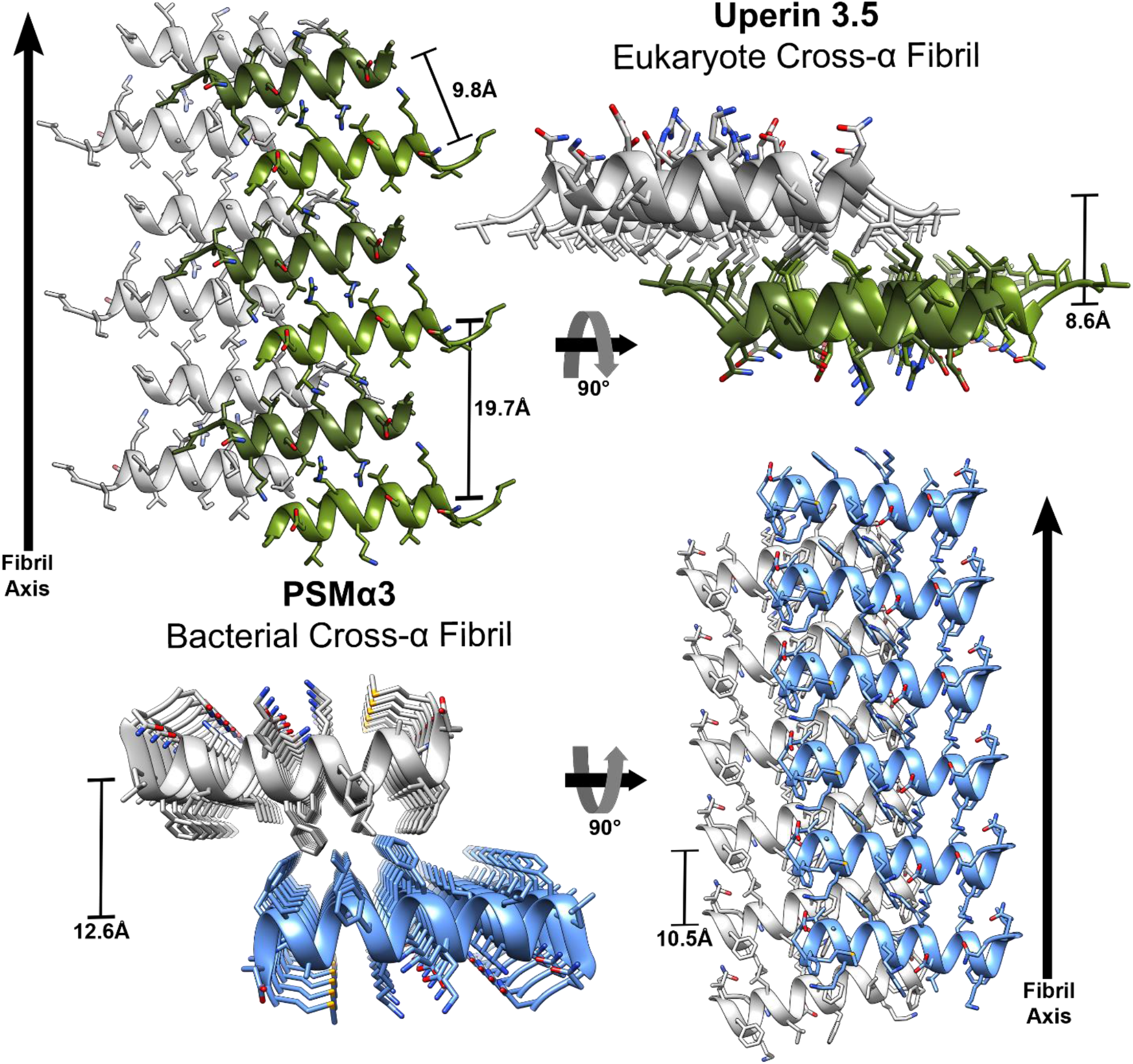
The cross-α amyloid fibril architectures of uperin 3.5 versus PSMα3. The crystal structures of uperin 3.5 (PDB ID: 6GS3) (green and grey ribbons) and PSMα3 (PDB ID: 5I55)^30^ (blue and grey ribbons) cross-α fibrils are shown in two orientations: down and along the fibril axis. Two mated helical sheets are displayed in each panel, with six layers of laterally stacked α-helices depicted in each sheet (fibrils are likely composed of thousands of layers). The α-helical sheets interact via their hydrophobic face to create a tight interface. In contrast to the parallel orientation of the helices along the sheets of PSMα3, the helices in uperin 3.5, colored in different shades of green for clarity, take on an antiparallel orientation. The distances between helices along the sheets and between sheets are shown in each panel. In both panels, side chains are shown as sticks with heteroatoms colored by atom type (nitrogen in blue, oxygen in red and sulfur in yellow).

While the antibacterial amphibian uperin 3.5 and the cytotoxic bacterial PSMα3 showed low sequence similarity (Figure S2A), both contained amphipathic helices and assembled into helical sheets that mate via a hydrophobic core (shown for uperin 3.5 in Figure S2 and Table S1). However, while the PSMα3 helices are orientated in a parallel fashion^30^, uperin 3.5 helices were arranged in an antiparallel manner. Thus, the cross-α amyloid fibril architecture can be encompassed by either parallel or antiparallel helical sheets. Uperin 3.5, in comparison to PSMα3, showed shorter inter-sheet and inter-helix distances, likely due to less bulky side chains (Figure 1). The high-order crystal packing of uperin 3.5 showed alternating hydrophobic and hydrophilic interfaces between rows of sheets (Figure S3), as seen in PSMα3^30^. The solvent-accessible surface area buried within the mated sheets of the cross-α uperin 3.5 fibril resembled that of PSMα3 (Table S1), despite the shorter sequence of uperin 3.5. Nevertheless, the shape complementarity between the uperin 3.5 mated sheets was smaller than that of mated PSMα3 sheets (Table S1), which might be related to the staggered orientation of the sheets (Figure 1). The staggered orientation might be compensated by higher-order packing in the fibril, with more than one pair of sheets per row (Figure S3). Overall, the eukaryotic uperin 3.5 and the bacterial PSMα3 share a cross-α architecture but display extensive polymorphism in helix orientation and stacking.

In addition to the extensive hydrophobic core between sheets along the fibril, uperin 3.5 was also stabilized via an array of inter-helical polar bonds that ran along each sheet. These included inter-helical electrostatic interactions between Asp4 and Lys14, and an inter-helical hydrogen bond between the side chain of Arg7 and the amide carbonyl of Arg7 from an adjacent helix (Figure S2D). Overall, each helix is potentially involved in four polar interactions along the sheet of stacked helices forming the cross-α fibril. Previous studies have indicated the importance of Arg7 in lipid interactions^40,41^, which aligns with our hypothesis, discussed below, that membrane interactions and the formation of cross-α fibrils are correlated processes.

### Uperin 3.5 shows a chameleon propensity of a secondary structure switch in the presence of bacterial membrane lipids

In solution, uperin 3.5 formed a random-coil structure with typical minima at 197 nm, as measured using circular dichroism (CD) spectroscopy. This was in contrast to PSMα3, which is helical in solution^31^. Nevertheless, the addition of small unilamellar vesicles (SUVs) composed of 1,2-dioleoyl-sn-glycero-3-phosphoethanolamine (DOPE) and 1,2-dioleoyl-sn-glycero-3-phospho-(1’-rac-glycerol) (DOPG) at a 1:3 molar ratio, mimicking a Gram-positive bacterial membrane^43,44^, induced an immediate secondary structure transition of uperin 3.5 towards an α-helical conformation, with typical minima near 208 and 222 nm (Figure 2A). A similar phenomenon has been reported for other AMPs, which showed a random coil structure in solution, and a structural transition in the presence of membrane lipids^45^.

**Figure 2.**
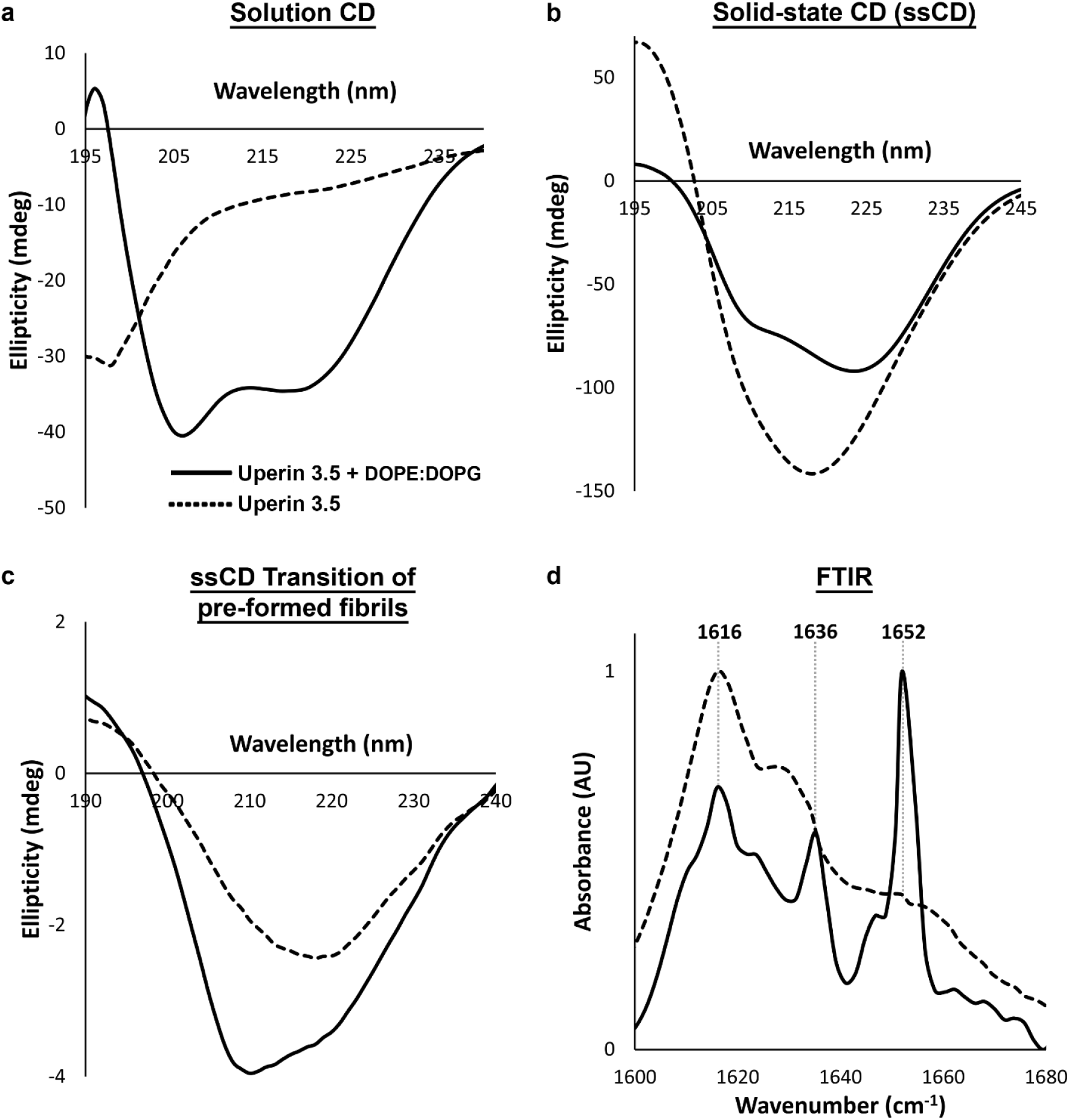
Bacterial lipids induce uperin 3.5 secondary structure transition into α-helical species in soluble and fibrillar states. (**a**) Solution CD spectra of uperin 3.5, indicating a random coil conformation (dashed curve). Upon addition of DOPE:DOPG SUVs, the uperin 3.5 structure immediately transitioned and stabilized in a dominant α-helical conformation (solid curve). (**b**) Solid-state CD (ssCD) spectra of uperin 3.5 thin films, indicating the formation of β-rich fibrils (dashed curve). When incubated with DOPE:DOPG SUVs, the fibrils took on a α-helical conformation (solid curve). The ssCD spectrum of the cross-α fibrils of PSMα3 are shown in Figure S4 for comparison. (**c**) ssCD spectra of an uperin 3.5 thin-film, indicating the formation of β-rich fibrils (dashed curve). The addition of DOPE:DOPG SUVs solution on to the thin film of pre-formed β-rich fibrils resulted in a secondary structure transition towards the α-helical conformation (solid curve). (**d**) ATR-FTIR spectra of the amide I’ region of uperin 3.5 fibrils (dashed curve) showed a major peak at 1616 cm^−1^, indicative of cross-β amyloids^56–58^. Uperin 3.5 fibrils formed in the presence of DOPE:DOPG SUVs (solid curve) demonstrated a major peak at 1652 cm^−1^, indicative of α-helices ^59,60^ and a minor peak at 1616 cm^−1^, indicative of residual β-rich fibrils. The dotted grey lines indicate wavenumbers of 1616, 1634 and 1652 cm^−1^. In all experiments (a-d), the signal of buffer only or DOPE:DOPG SUVs solution was negligible and subtracted from each measured sample.

The secondary structure of dry uperin 3.5 fibrils was evaluated using solid-state CD (ssCD) spectroscopy^46–50^, previously shown to be highly applicable for characterization of aggregating proteins and peptides^47,51–53^. The ssCD spectrum measured for uperin 3.5 incubated in the absence of bacterial lipids indicated a β-rich conformation, with a typical minimum at 218 nm (Figure 2B). In contrast, uperin 3.5 fibrils incubated in the presence of DOPE:DOPG SUVs had a seemingly α-helical conformation (Figure 2B). While the fibril sample in the ssCD spectrum showed a deeper 222 nm minimum as compared to the 208 nm minimum, the sample in solution showed the opposite trend, with a deeper 208 nm minimum (Figure 2). The same trend was observed when analyzing a thin film of pre-formed PSMα3 fibrils (Figure S4). The switch in the depth of the minima of fibrils versus soluble uperin 3.5 could be related to different secondary structure subpopulations, association between α-helices^54,55^, or other physicochemical properties. Adding DOPE:DOPG SUVs to the pre-formed β-rich uperin 3.5 fibrils, led to a transition towards an α-helical conformation (Figure 2C), providing further evidence of lipid-induced helicity, even after the fibrils were already formed.

The secondary structure of dry uperin 3.5 fibrils was further studied using attenuated total internal reflection Fourier-transform infrared (ATR-FTIR) spectroscopy, which was shown useful for characterization of fibrillar architectures in amyloid proteins and peptides^56–60^. The major peak in the amide-I’ region of the IR absorption spectrum of uperin 3.5 fibrils was at 1616 cm^−1^, indicative of the cross-β amyloid architecture^56–58^, while the minor peak at 1652 cm^−1^, was indicative of α-helices^59,60^ (Figure 2D), suggesting a mixed population with a predominant cross-β structure. Forming the fibrils in the presence of DOPE:DOPG SUVs resulted in a shifted IR spectrum, with a strong major peak at 1652 cm^−1^, indicative of a majority of α-helices (Figure 2D). Additional minor peaks at 1616 and 1634 cm^−1^, indicative of cross-β amyloids and regular β-sheets, respectively, suggested some remaining β-rich structures. The X-ray fiber diffraction of uperin 3.5 incubated in the absence of bacterial lipids, showed a cross-β signature^61^, with orthogonal reflection arcs at 4.7 Å spacing (sharp) and at 7-11 Å (diffused) (Figure S5). A fiber X-ray diffraction of uperin 3.5 fibrils grown in the presence of bacterial lipids showed strong reflections of the lipids, and the pattern of the protein was not adequately coherent for interpretations. Overall, these findings suggest a chameleon behavior of uperin 3.5, with transition from a random coil conformation in its soluble state to either a cross-β fibril conformation, formed in lipid-free solution, or to cross-α fibril structure, induced by the presence of bacterial membrane lipids.

### Thermostability of uperin 3.5 fibrils might stem from their chameleon propensity

Uperin 3.5 fibrils were found to be thermostable, as observed in electron micrographs after a heat shock treatment of incubating the fibrils at 60°C for 10 min (Figure 3B). The ssCD spectrum of preformed uperin 3.5 fibrils indicated largely on β-rich species (Figure 2C&3B), while after the heat shock treatment a deeper minimum at 218 nm was observed, as compared with non-heat-shock treated fibrils of the same sample, indicating a larger population of β-rich species (Figure 3A). In comparison, *S. aureus* PSMα1 fibrils which bear a cross-β configuration^29^, were also thermostable, while the PSMα3 cross-α fibrils dissolved after the heat-shock treatment (Figure S6), indicating on a fibril secondary-structure dependent thermostability.

**Figure 3.**
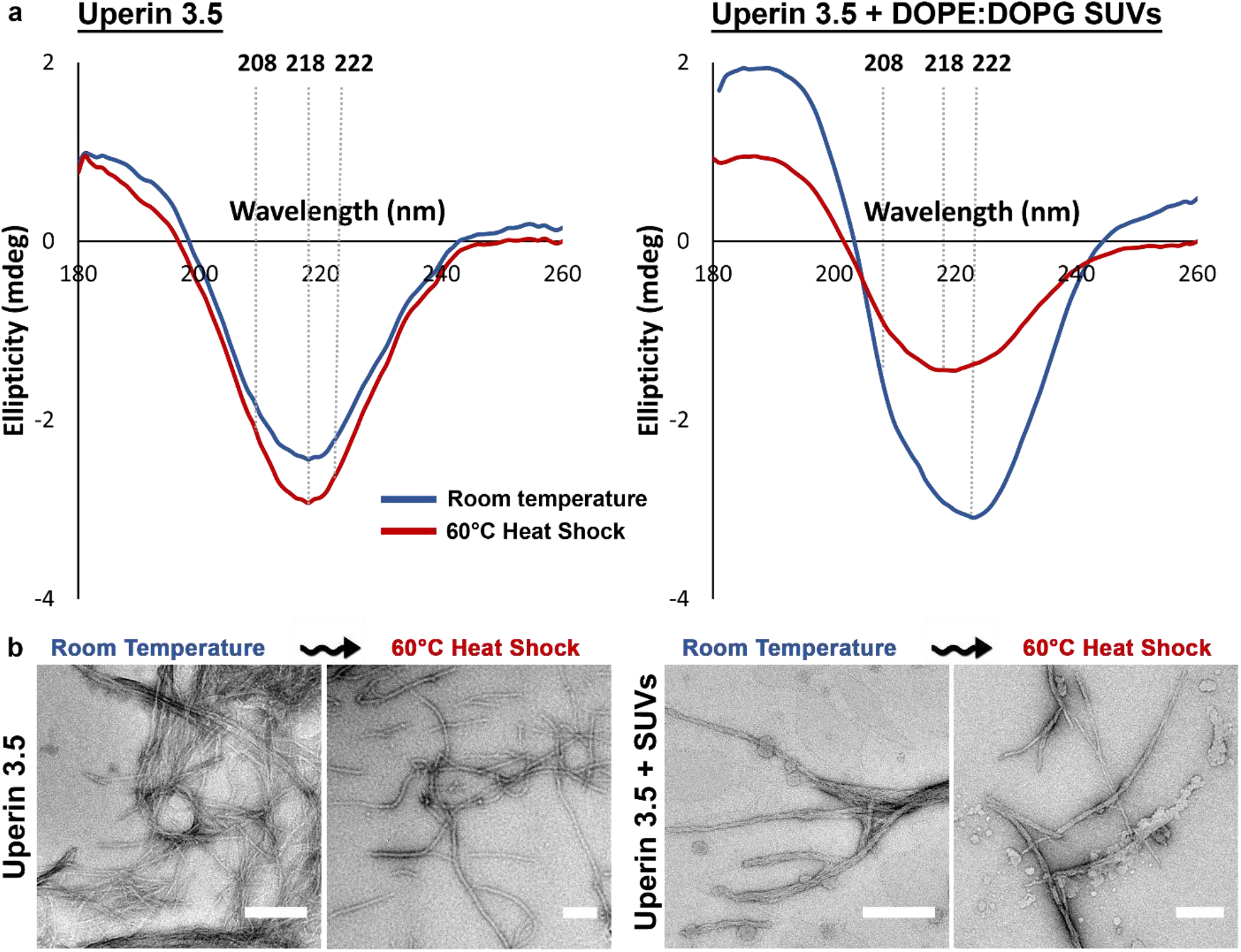
Thermal stability and the effect of heat on the secondary structure of uperin 3.5 fibrils. (**a**) Solid-state CD (ssCD) spectra showing the effect of heating on the secondary structure of preformed thin films of uperin 3.5 fibrils formed alone or in the presence of DOPE:DOPG SUVs. Left panel: uperin 3.5 incubated alone showed a predominant β-rich fibril conformation before and after a 10 min, 60°C heat shock treatment (blue and red curves, respectively). Right panel: uperin 3.5 fibrils formed in the presence of DOPE:DOPG SUVs showed predominant α-helical fibril conformation (blue curve), while after a 10 min, 60°C heat-shock treatment, it showed a secondary structure transition towards a β-rich conformation (red curve). The dotted grey lines indicate wavelengths of 208 nm, 218 nm, and 222 nm. The same sample was tested following temperature change in both assays. (**b**) TEM micrographs of 1 mM uperin 3.5 incubated for 2 days at room-temperature alone, or in the presence of DOPE:DOPG SUVs, showing thermostable fibrils. Scale bars represent 300 nm. The same sample was tested following temperature change in both assays.

Uperin 3.5 incubated in the presence of DOPE:DOPG SUVs showed massive fibril formation and thick fibrils around and on the SUVs (Figure 3B). The ssCD spectra indicated on a mixed population of species with a predominant α-helical conformation (Figure 3A). When subjected to a heat shock treatment, the uperin 3.5 fibrils formed in the presence of the SUVs remained stable, whereas the SUVs seemed to partially denature, as observed in the electron micrographs (Figure 3B), and the ssCD spectra indicated on a transitioned to a predominant β-rich architecture (Figure 3A). Overall, these findings demonstrate a heat-induced transition of uperin 3.5 fibrils to a predominantly β-rich conformation, which probably led to thermostability. This corresponds to the higher stability of the cross-β PSMα1 as compared to cross-α PSMα3 (Figure S6), which are homologous sequences with fibrils of different secondary structures.

### Uperin 3.5 fibrillation is involved in antibacterial activity

The antibacterial activity of uperin 3.5, examined using the disc (agar) diffusion test against four Gram-positive pathogens, demonstrated its more potent activity against *Micrococcus luteus* as compared to *Staphylococcus hominis*, *Staphylococcus epidermidis* and *Staphylococcus aureus* (Figure S7), as previously reported^38^. Using the standard broth dilution assay, a minimum inhibitory concentration (MIC) of freshly dissolved uperin 3.5 against *M. luteus*, defined as the lowest concentration that prevented bacterial growth for 24 h, was 2 μM (Table 2). Membrane surface topography analysis of *M. luteus* bacterial cells, visualized using scanning electron microscopy (SEM), indeed showed that overnight incubation with 2 μM uperin 3.5 induced some membrane damage, while 6 μM uperin 3.5 induced severe morphological damage (Figure 4A). Higher-resolution TEM micrographs of 4 μM uperin 3.5 incubated for 24 h with *M. luteus* showed massive fibril formation around the bacterial cells, which led to cell death (Figure 4B). Uperin 3.5 incubated under the same conditions without the bacteria showed only scarce fibril formation (Figure 4B), suggesting facilitated aggregation in the presentence of bacterial cells. A similar effect was observed in samples of uperin 3.5 with DOPE:DOPG SUVs, in which massive fibril formation was observed around the SUVs (Figure 3A).

**Table 2.**
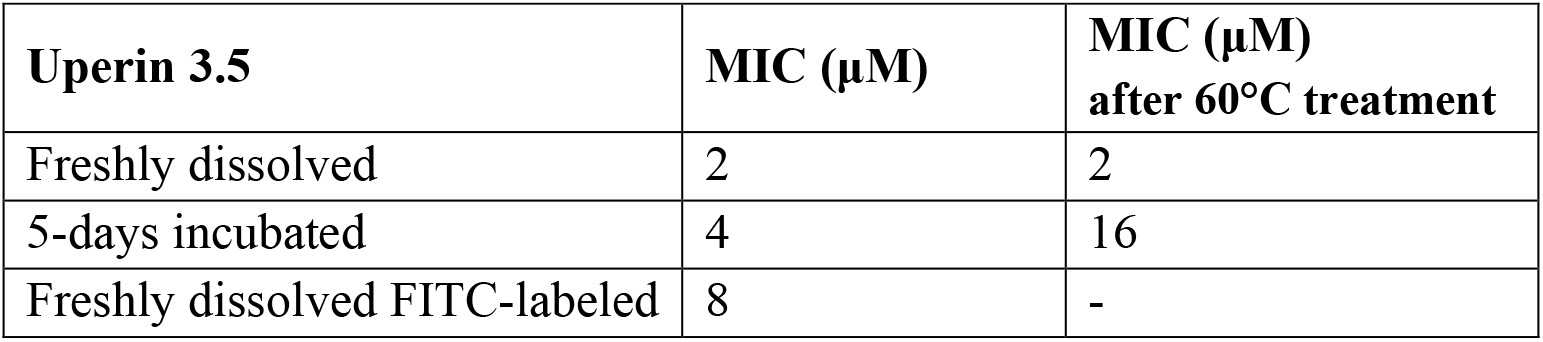
Uperin 3.5 antibacterial activity against *Micrococcus luteus*.

**Figure 4.**
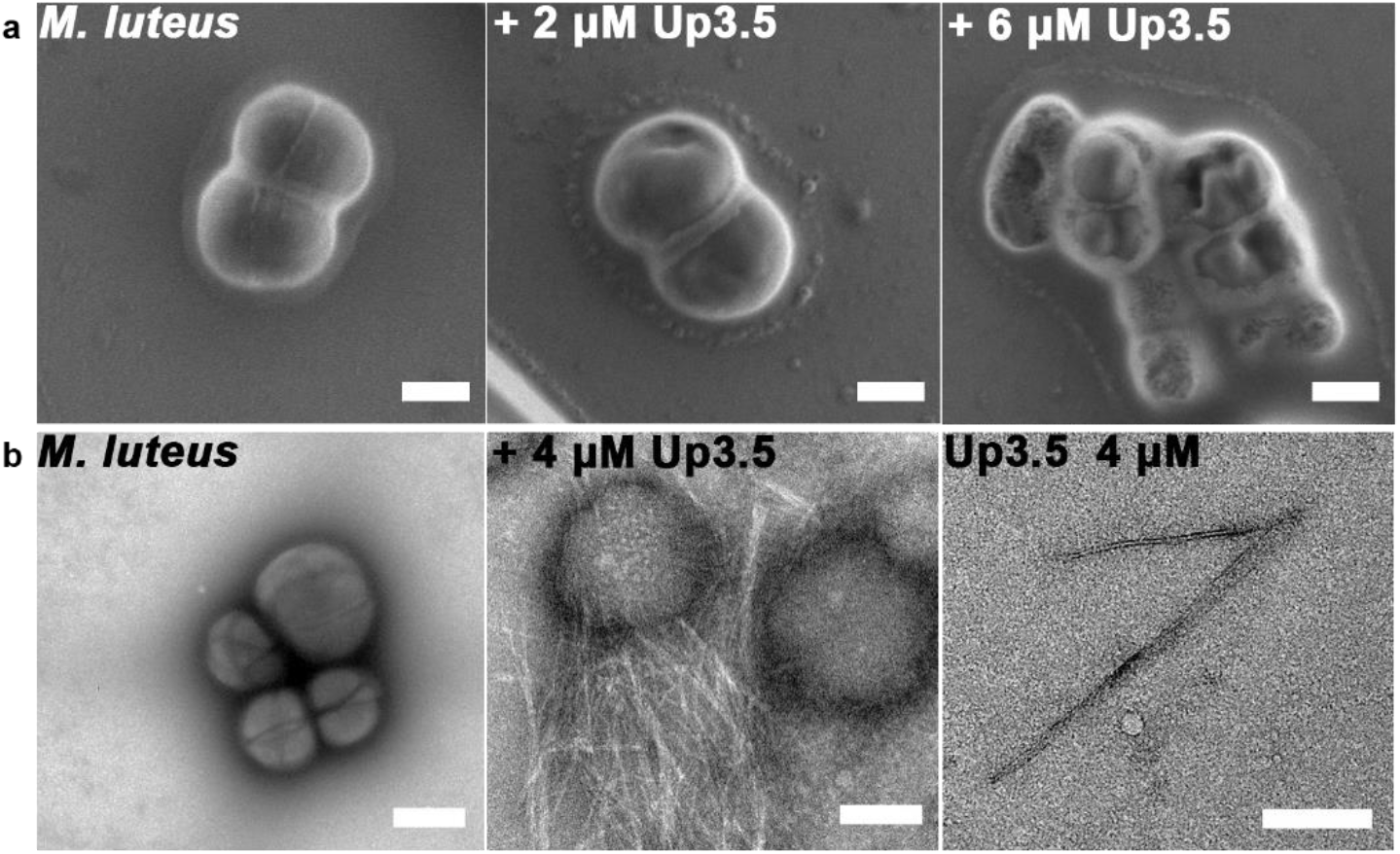
Electron micrographs of uperin 3.5 fibrils formed on bacterial cells and of uperin 3.5-induced membrane damage. (**a**) Scanning electron microscopy (SEM) images showing *M. luteus* cells incubated for 24 h in the absence (left) or presence of uperin 3.5 at 2 μM (middle) and 6 μM (right), and resultant membrane dents. Scale bars represent 500 nm for all images. (**b**) Transmission electron micrographs of *M. luteus* cells incubated in the absence (left) or presence (middle) of 4 μM uperin 3.5 for 24 h, showing massive fibril formation around bacterial cells (middle panel). Scarce fibril formation is shown upon incubation without cells (middle panel). Scale bars represent 1 μm for *M. luteus* (left), and 200 nm for *M. luteus* incubated with uperin 3.5 (middle) and for uperin 3.5 incubated alone (right).

In accordance with the above findings, light microscopy images showed that FITC-labelled uperin 3.5, which retained fibril-forming and antibacterial abilities (Figure S8 and Table 2), accumulated on or inside rhodamine-labelled DOPE:DOPG SUVs (Figure S9A) or *M. luteus* cells (Figure S9B). The latter led to membrane permeation, followed by propidium iodide uptake, indicating cell death. Taken together, these findings demonstrate the bi-directional effects of uperin 3.5 and bacterial cells, namely the role of bacterial lipids in inducing cross-α fibrillation, and the role of cross-α fibrillation in antibacterial activity.

The heat shock-treated uperin 3.5 soluble peptide retained its antibacterial function, with a similar MIC against *M. luteus* cells, before as compared to after heat treatment (Table 2). In contrast, pre-formed uperin 3.5 fibrils (pre-incubated for 5 days) exhibited a heat-induced reduction in their antibacterial activity, with a MIC of 16 μM versus 4 μM following as compared to before heat treatment, respectively (Table 2). Of note, it is difficult to compare the activity of fresh and incubated samples due to the rapid aggregation rate and change in effective concentration. The heat-induced effect was compared using the same sample immediately pre-and post-heating, yet potential changes in fibril dynamics must also be considered when analyzing the factors that affect activity. Moreover, since heat and the presence of lipids had opposite effects on the secondary structure of pre-formed uperin 3.5 fibrils (Figures 2&3A), it must be assumed that addition of uperin 3.5 to bacterial cells may partially reverse some of the heat-induced effects. Thus, the overall four-fold reduction in antibacterial activity of pre-formed fibrils after heating is likely the result of a combined effects of heat-induced secondary structure transition towards a β-rich conformation, altered dynamics between fibrils and monomers, changes in biophysical properties and effective concentration, and additional factors.

## Discussion

Here, we revealed and characterized the functional cross-α fibril architecture formed by the eukaryotic uperin 3.5 AMP, and its chameleon secondary structure switch, putatively regulating activity. The helical fibril formation of uperin 3.5 was largely induced by, and formed on bacterial cells, where it consequently led to severe membrane damage and cell death (Figures 2,4&S9). Of note, crystal structures of short helical AMPs are rather rare^62^, nevertheless, quasiracemate crystal structures of frog AMP magainin 2 derivatives showed laterally stacked helices^63^, similarly to the cross-α architecture. While correlations between amyloid formation and antibacterial activity have long been drawn^6–16,20–26,64^, this work provided an atomic-level description for this link, involving a cross-α amyloid configuration. The departure of the secondary structure of the cross-α amyloid uperin 3.5 from the typical cross-β configuration clarifies the connection between AMPs which are largely helical in nature, and amyloid formation. Interestingly, amyloids secreted from microbes, either during a pathogenic infection, or even by the gut microflora, have been implicated in the development of neurodegenerative and amyloid systemic disorders^65,66^. The overall connection between microbes, AMPs, the immune system, and neurodegenerative and systemic diseases requires more extensive studies.

In line with the reported polymorphism of cross-β amyloids^67^, we observed polymorphism within the cross-α fibrils of uperin 3.5 and PSMα3. Both showed helical mated sheets of stacked amphipathic α-helices, which lay perpendicular to the fibril axis. PSMα3 contained parallel α-helices, while uperin 3.5 contained antiparallel helices within the sheets (Figure 1), further recapitulating the parallel and antiparallel β-sheet configurations. Similarly, an antiparallel cross-α crystal structure was shown in a designed synthetic peptide^34^. Furthermore, the inter-helix and inter-sheet distances of uperin 3.5 were smaller than those in PSMα3, likely due to less bulky side chains, and due to the staggered sheets of uperin 3.5 (Figure 1). As amyloid polymorphism has been correlated with different toxicity level and prion disease strains^68,69^, the polymorphisms within cross-α structures may dictate toxicity levels, mechanisms of action, cell specificity, and fibril stability. While PSMα3 was stable as an α-helix in solution and in fibrils, uperin 3.5 helicity was dependent on the presence of lipids or other components. Moreover, while PSMα3 cross-α fibrils dissolved when subjected to heat shock, uperin 3.5 was thermostable (Figures S6 and 3B), which is likely attributed to the heat-induced transition into cross-β fibrils (Figure 3A). In PSMα3, secondary structure changes were only inducible via mutations^31^ or truncations^29^ which formed mixed α/β populations or exclusively β-rich structures^31^.

Chameleon proteins that can adopt diverse secondary structures in response to external chemical or physical stimuli have been previously described^70–72^, some induced by lipids or heat,^73–75^, similar to what was observed here for uperin 3.5 fibrils (Figures 2 and 3). Martin, Carver and co-workers previously pointed on a potential chameleon propensity of uperin 3.5^41^, and discussed the role of helices as intermediates to aggregation^33,41,76,77^ along with the potential contribution of intermolecular helix–helix interactions in the formation of a fibrillation nuclei^41^. A thermal-induced cross-α/cross-β transition was previously reported in a synthetic self-assembling heptapeptide, suggested to occur via an intermediate turn motif^33^. In addition, a dipeptide protein-mimics, containing triazole linkages as peptide-bond surrogate, formed β-sheet rich structures in crystals, while heat-induced polymerization into a pseudoprotein revealed transition into helical sheets resembling the cross-α configuration^37^. The seemingly relatively unstructured C-terminal region of uperin 3.5 (Figure 1), might be essential for the mechanism of cross-α/cross-β transition. An alternative suggestion for the transition mechanism of uperin 3.5 could stem from environmental effects on the equilibrium of mixed α/β species (Figure 2).

Taken together, we hypothesize that in the amphibian uperin 3.5, secreted on the skin of the toadlet^38^, the chameleon secondary structure switch could be related to regulation of activity. Namely, uperin 3.5 monomers secreted on the amphibian skin have disordered structures, while the presence of bacterial cells induces helicity and cross-α formation, which putatively facilitates the antibacterial activity and eliminates the threat of infection. In the absence of bacteria, uperin 3.5 assembles into inactive cross-β fibrils. This is somewhat similar to human amyloid toxicity which is debatably attributed to prefibrillar oligomeric conformations, which can contain α-helices^76^, while the mature β-rich fibrils are considered less toxic^78^. The β-rich uperin 3.5 can also serve as a reservoir^33,79^, presenting a means of storing a high local concentration of inactive uperin 3.5 on the amphibian skin, which can be switched back to helical conformation with the appearance of bacterial cells (as tested on mimetics, Figure 2C). We expect that additional studies will shed additional light on the underlying regulation.

The mechanism by which uperin 3.5 disrupts membranes is yet to be fully determined. Recently, we suggested a peptide-membrane co-aggregation model for PSMα3 toxicity against human cells^31^. In this model, toxicity is not caused by a particular entity, such as monomers, oligomers or fibrils, but, rather, entails a dynamic process of peptide aggregation that is induced by, and involves, membrane lipids^31^. Thus, both the presence of soluble species and the ability to fibrillate are critical determinants of toxicity. We suggest similar aggregation dynamics for uperin 3.5, in which cross-α fibrillation is both directly related to the presence of, and incorporates, membrane lipids (Figure S9). However, we expect differences in the mechanisms of action of PSMα3 and uperin 3.5, due to the different properties of their fibrils and different sequences.

Altogether, the presented findings emphasize the structural complexity and the polymorphic nature of the amyloid fold, as shown in cross-α fibrils formed by prokaryotic and eukaryotic toxic peptides, namely the bacterial PSMα3 and the amphibian uperin 3.5. This extends the definition of amyloid structures, and of functional fibrils. The findings could lay the foundations for the design of novel synthetic AMPs with enhanced potency, stability, and bioavailability, and with controllable storage and activities, to fight the growing threat of aggressive and resistant microbial infections.

## Materials and Methods

### Peptides and reagents

Uperin 3.5 peptide (Uniprot accession code: P82042) of the sequence GVGDLIRKAVSVIKNIV-NH2, fluorescein isothiocyanate (FITC)-labeled uperin 3.5 of the sequence GVGDLIRKAVSVIKNIV-K(FITC)-NH_2_, PSMα1 (Uniprot accession code: H9BRQ5) of the sequence MGIIAGIIKVIKSLIEQFTGK and PSMα3 (Uniprot accession code: H9BRQ7) of the sequence MEFVAKLFKFFKDLLGKFLVNN (all custom synthesis) at >98% purity were purchased from GL Biochem Ltd. (Shanghai, China) or GenScript (USA). Amidation of the C-termini of AMPs occurs naturally, decreasing the negative charge of the free carboxyl end of the peptide; therefore, uperin 3.5 was synthesized with an amidated C-terminal or with unmodified termini for crystallography. FITC labeling is mediated by the charged epsilon amine on lysine residues. In order to preserve the propensity of uperin 3.5 aggregation, and its overall charge, FITC was conjugated to an additional lysine residue, added at the C-terminus. Dimethyl sulfoxide (DMSO), chloroform, methanol, acetone, hydrochloric acid (HCl), 1,1,1,3,3,3-hexafluoro-2-propanol (HFIP), thioflavin T (ThT), deuterium oxide (D_2_O), uranyl acetate, poly-L-lysine, and propidium iodide were purchased from Sigma-Aldrich. Sodium chloride, potassium dihydrogen phosphate and potassium phosphate dibasic were purchased from Merck. Ultra-pure water was purchased from Biological Industries (Israel). Luria-Bertani (LB) and brain heart infusion (BHI) bacterial media were purchased from Difco-BD, USA. Lipids, including 1,2-dioleoyl-sn-glycero-3-phosphoethanolamine (18:1 Δ9-Cis DOPE), 1,2-dioleoyl-sn-glycero-3-phosphoethanolamine-N-(lissamine rhodamine B sulfonyl 18:1 Δ9-Cis DOPE), and 1,2-dioleoyl-sn-glycero-3-phospho-(1’-rac-glycerol) (18:1 Δ9-Cis DOPG) were purchased from Avanti Polar Lipids, Inc. (USA).

### Peptide pre-treatment

Lyophilized synthetic uperin 3.5 and FITC-labelled uperin 3.5 were dissolved to a concentration of 1-5 mg/ml in HFIP, and then sonicated for 10 min, in a bath sonicator, at room temperature. The organic solvent was evaporated using a mini rotational vacuum concentrator (RVC 2-18 CDplus Christ, Germany) at 1,000 rpm, for at least 2 h at room temperature. Treated peptide was stored dry, at −20 °C, until use.

### Small unilamellar vesicle (SUV) preparation

Lipids were dissolved to 25 mg/ml stock solutions in chloroform and then mixed in the following combinations and ratios: DOPE:DOPG 1:1, DOPE:Rhod-DOPE:DOPG 0.23:0.08:0.31. The solvent was then evaporated under vacuum to complete dryness. The lipid film was rehydrated in a solution containing 50 mM potassium phosphate buffer and 150 mM sodium chloride, pH 7.3, vortexed and incubated for several hours to overnight. Final lipid concentration was 10 mM. The lipid suspension was sonicated, on ice, using a tip sonicator (SONICS, USA), at 20 % amplitude, for 5 min, with 10 sec pulses, to form SUVs. The solution was incubated (1 h) at room temperature, to stabilize prior to use.

### Fibrillation assays

#### Thioflavin T (ThT) fluorescence fibrillation kinetics assay

Thioflavin T (ThT) is a widely used “gold standard” stain for identifying and exploring formation kinetics of amyloid fibrils in vitro^83^. Fibrillation curves in the presence of ThT commonly show a lag time for the nucleation step, followed by rapid aggregation. Pre-treated uperin 3.5 peptide aliquots were re-dissolved in ultra-pure water to a 10 mM stock solution, sonicated in a bath sonicator for 5 min, at room-temperature (RT), and immediately diluted to 1 mM in a 20 mM potassium phosphate buffer and 100 mM NaCl, at pH 7.3. The reaction samples contained filtered ThT diluted from a 2 mM stock made in ultra-pure water. Final concentrations for each reaction were 100 μM peptide and 20 μM ThT. Blank samples contained everything but the peptide. The reaction was carried out in a black 96-well flat-bottom plate (Greiner Bio-One), covered with a thermal seal film (EXCEL Scientific, USA) and incubated in a plate reader (CLARIOstar or FLUOstar Omega, BMG LABTECH, Germany), at 37 °C, with 500 rpm shaking, for 85 sec, before each reading cycle, and up to 1000 cycles of 6 min each. Measurements were made in triplicates. Fluorescence was measured at an excitation wavelength of 438±20 nm and emission of 490±20 nm, over a period approximately 100 h. All triplicate values were averaged, blank readings were subtracted from all other readings, and the results were plotted against time. Calculated standard errors of the mean are presented as error bars. The entire experiment was repeated at least three times, on different days.

#### Transmission electron microscopy (TEM)

Sample preparation before fixation on TEM grids are described here for each figure separately. Figure S1: Uperin 3.5 (100 μM) was collected following the ThT fibrillation kinetics assay (described above) by combining the contents of 2-3 wells from the plate. The samples were centrifuged at 21,000 g and the supernatant was discarded. The pellet containing the fibrils was resuspended in 20 μl ultra-pure water. Figure 5: To visualize uperin 3.5 fibrils formed in the presence of bacterial cells, 4 μM uperin 3.5 was incubated with *M. luteus* for 24 h, at 30 °C. After testing the antimicrobial activity using broth-dilution assay (described below), the sample analyzed using TEM. A sample of uperin 3.5 incubated without bacterial cells, under the same conditions, served as a control.

Samples (5 μl) were applied directly onto 400-mesh copper TEM grids with Formvar/Carbon support films (Ted Pella), which were glow-discharged (PELCO easiGlow, Ted Pella) immediately before use. Samples were allowed to adhere for 2 min and negatively stained with 5 μl 2 % uranyl acetate solution. Micrographs were recorded using a FEI Tecnai G2 T20 S-Twin transmission electron microscope, at an accelerating voltage of 200 KeV (located at the Department of Materials Science and Engineering, Technion, Israel). Images were recorded digitally with a Gatan US 1000 CCD camera, using the Digital Micrograph® software.

##### Thermostability assays (figures 3 and S6)

Uperin 3.5 and PSMα3 were dissolved to 10 mM in ultra-pure water, and PSMα1 was dissolved to 10 mM in DMSO. Uperin 3.5, PSMα1 and PSMα3 were further diluted to 1 mM, while a portion of the 10 mM sample of uperin 3.5 was diluted into a DOPE:DOPG SUVs solution (prepared as described above) to final concentrations of 1mM uperin 3.5 and 3 mM SUVs (1:3 peptide:lipid molar ratio). Samples were incubated at room temperature for two days and then applied directly onto copper TEM grids and stained, as described above. The samples were then incubated at 60 °C in a pre-heated thermoblock, for 10 min, and immediately applied onto TEM grids and stained. Micrographs were recorded as described above, and in the case of too thick samples, they were re-prepared similarly but diluted 10-fold.

### Crystallization conditions

Non-pretreated uperin 3.5 peptide synthesized with free (unmodified) termini, was used for crystallization experiments. The peptide was dissolved in ultra-pure water, to 10 mM, followed by a 10 min sonication in a bath sonicator, at room temperature. Peptide solution drops (100 nl) were dispensed by the Mosquito automated liquid dispensing robot (TTP Labtech, UK), onto crystallization screening plates. All crystals were grown, at 20 °C, via hanging-drop vapor diffusion. The drop was a mixture of peptide in reservoir solution (0.1 M KSCN, 0.1 M MES pH 6.03 and 20 %v/v Jeff 600). Crystals grew after a few days and were flash-frozen with cryoprotection of 20 % ethylene-glycol and stored in liquid nitrogen prior to data collection.

### Structure determination and refinement

X-ray diffraction data were collected at 100 K, at the micro-focus beamline Massif-III (ID30a-III) of the European Synchrotron Radiation Facility (ESRF) in Grenoble, France; wavelength of data collection was 0.9677 Å. Data indexation, integration and scaling were performed using XDS/XSCALE^84^. Phases were obtained using ARCIMBOLDO_LITE^85^, which uses PHASER to place individual α-helices by eLLG-guided molecular replacement^86^, then expands partial solutions with SHELEX^87^, through density modification and autotracing, into a complete model. Model building was done using Coot^88^. Crystallographic refinements were performed with Refmac5^89^, and illustrated with Chimera^90^. There were no residues that fell in the disallowed region of the Ramachandran plot. Crystallographic parameters are listed in Table 1.

#### Calculations of structural properties

*Hydrophobicity:* The hydrophobicity scale presented in the figures was created using Chimera^90^, which is based on the hydrophobicity scale of Kyte and Doolittle^91^. *Inter-sheet distance:* The amphipathic nature of the peptides results in alternating rows of dry and wet interfaces in the crystal packing. The dry interface corresponds to the hydrophobic packing between mated sheets. In the wet interface, interactions between sheets are mediated by water molecules. The inter-sheet and inter-helix distances were calculated in Chimera^90^. Of note, these calculations were performed using a different method than in our first report on PSMα3^30^, therefore resulting in a slightly different inter-sheet value for the WT PSMα3. *Shape complementarity:* The Lawrence and Colman’s shape complementarity index^92^ was used to calculate shape complementarity between pairs of sheets forming the dry interface. *Area buried within the dry interface:* Area buried was calculated using AREAIMOL^93,94^ with a probe radius of 1.4Å. Calculations were performed using the CCP4 package^95^. The area buried is the sum of the differences between the accessible surface areas of a single molecule and when it is in contact with the other helices on the same sheet or opposite sheets, as indicated in Table S1.

### Solution circular dichroism (CD) spectroscopy

Immediately prior to CD experiments, pre-treated uperin 3.5 was dissolved to 10 mM in ultra-pure water and diluted to 1 mM in 20 mM potassium phosphate buffer and 100 mM NaCl, pH 7.3. The signal from blank solutions of either buffer alone or of buffer with 0.6 mM SUVs composed of DOPE:DOPG, were recorded just before the addition of uperin 3.5 to the cuvette. Uperin 3.5 was diluted in the CD cuvette to a final concentration of 0.2 mM, in the presence or absence of 0.6 mM SUVs. Far-UV CD spectra were recorded with an Applied Photophysics PiStar CD spectrometer (Surrey, UK), using a 1 mm path-length quartz cell (Starna Scientific, UK), at a temperature of 25 °C. Changes in ellipticity were monitored from 260 nm to 180 nm, at 1 nm step sizes and a bandwidth of 1 nm. The measurements shown are an average of three scans for each sample or two scans for blanks, captured at a scan rate of 1 sec per point, with appropriate blanks subtracted. Regardless, the blanks of buffer or DOPE:DOPG SUVs solution had a negligible signal. The CD values in Figure 2A are shown as ellipticity (millidegrees) for sake of comparison with the solid-state CD spectra.

### Solid-state circular dichroism (ssCD) spectroscopy

Uperin 3.5 (5 mg/ml) dissolved in ultra-pure water and uperin 3.5 (1.8 mg/ml) mixed with DOPE:DOPG SUVs solution at a 1:3 peptide:lipid molar ratio were incubated at 37 °C, with 300 rpm shaking, for 2 days. PSMα3 (5 mg/ml) dissolved in ultra-pure water was incubated at 37 °C, with 300 rpm shaking, for 2 days. Fibrillated samples were centrifuged at 10,000 g for 5 min and re-suspended in ultra-pure water. A small sample (30-40 μl) containing fibrils was then spread on a 13 mm fused silica disc (Applied Photophysics, UK) and incubated on a thermoblock, at 38 °C, for at least 5 min, or until the water evaporated completely, resulting in a visible thin film of fibrils. The disc was mounted into the solid-state sample holder (Applied Photophysics, UK). Far-UV ssCD spectra were recorded with an Applied Photophysics Chirascan CD spectrometer (Surrey, UK), at room-temperature. Changes in ellipticity were monitored between 260 nm and 180 nm, with a 1 nm step size and a bandwidth of 1 nm. To minimize the effect of sample orientation, the ssCD spectra were recorded at eight or four intervals of 45 or 90 degrees, respectively. Prior to preparation of each thin film, ssCD spectrum of the empty disc or disc with appropriate blank solution was measured. Regardless, the blanks of buffer or DOPE:DOPG SUVs solution had a negligible signal. In between measurements, the disc was thoroughly washed with ultra-pure water, 5 mM HCl solution, 2% Hellmanex III detergent solution (Hellma Analytics, Germany) and finally with acetone, and dried with a hot air blower. The measurements shown are an average of two or three scans for each blank or sample at each orientation, captured at a scan rate of 1 sec per point, with sample-specific blanks subtracted. Since the ssCD measurement is performed on a thin film of dehydrated fibrils, the exact concentration of the sample is unknown and the path length is influenced by the film’s thickness, both of which are crucial for calculating the molar ellipticity in order to determine the accurate percentage of secondary structures. Thus, ssCD data was interpreted according to the established spectra of secondary structures (e.g., a minimum at 218nm indicates a β-strand and double minima at ~208nm and ~222nm indicate a α-helix). For the sake of comparison, the solution CD spectrum shown in Figure 2A is also presented as ellipticity in millidegrees.

#### Thermostability assays (figure 4)

Uperin 3.5 pre-treated samples were dissolved in ultra-pure water to 1mM (1.8 mg/ml), mixed with DOPE:DOPG SUVs at a 1:3 peptide:lipid ratio and incubated for 5 days at room temperature. The ssCD spectra of each sample was measured as described above, at room temperature, and again after a 10 min, 60 °C heat-shock, while still being placed as a thin-film on the disc.

### Attenuated total internal reflections Fourier transform infrared (ATR-FTIR) spectroscopy

Uperin 3.5 was dissolved to 1 mg/ml in 5 mM hydrochloric acid (HCl) and sonicated in a bath sonicator for 5 min, at room temperature. The peptide solution was frozen in liquid nitrogen and lyophilized overnight to complete dryness. The procedure was repeated three times to completely remove trifluoroacetic acid (TFA) residues, as TFA has a strong FTIR signal at the amide I’ region of the spectra. SUVs composed of DOPE:DOPG were prepared as described above, with the exception of rehydration in D_2_O instead of H_2_O. Immediately prior to measurements, the dry peptide was dissolved either in D_2_O, with or without SUVs, to a final concentration of 20 mg/ml, with a 1:1 peptide:lipid molar ratio. Samples (5 μl) were spread on the surface of the ATR module and allowed to dry under nitrogen gas to purge water vapors. Uperin 3.5 without SUVs was analyzed using a MIRacle Diamond w/ZnSe lens 3-Reflection HATR Plate (Pike Technologies, USA), and uperin 3.5 with SUVs was analyzed using a Platinum ATR with a single-reflection diamond crystal (Bruker Optics). Absorption spectra were recorded on the dry samples using a LN-MCT detector on the Tensor 27 FTIR spectrometer (Bruker Optics). Measurements were performed in the wavelength range of 1500-1800 cm^−1^, at 2-4 cm^−1^ steps, and averaged over 1000 scans. Background (nitrogen gas) and blank (D_2_O or DOPE:DOPG SUVs in D_2_O) were measured and showed a negligible signal and subtracted from the final spectra. The data were edited using Origin Lab 2018 (OrigianLab Corporation, USA), wherein the blanks were subtracted, baseline was corrected, the spectra were normalized, and the second derivative was calculated in order to analyze peaks. The amide I’ region of the spectra (1600 – 1700 cm^−1^) is presented to show the relevant peaks (Figure 2D).

### Fibril X-ray diffraction

Uperin 3.5 peptide was re-dissolved in ultra-pure water to 20 mg/ml. Droplets (2 μl) were placed between two sealed-end glass capillaries and incubated until complete dryness (up to 1 h) at RT. X-ray diffraction of the fibrils was collected at the micro-focus beamline P14, operated by EMBL Hamburg, at the PETRA III storage ring (DESY, Hamburg, Germany) and at the ID29 beamline of the European Synchrotron Radiation Facility (ESRF) in Grenoble.

### Bacterial strains and culture media

*Micrococcus luteus* (an environmental isolate) was a kind gift from Prof. Charles Greenblatt from the Hebrew University in Jerusalem, Israel. *Staphylococcus hominis* (ATTC 27844), *Staphylococcus aureus* (ATCC 29213) and *Staphylococcus epidermidis* (ATCC 12228) were purchased. *M. luteus* was cultured in Luria-Bertani (LB) medium at 30°C, with 250 rpm shaking, overnight. *S. aureus, S. hominis* and *S. epidermidis* were cultured in brain-heart infusion (BHI) medium, at 37 °C, with 250 rpm shaking, overnight.

### Disc (agar) diffusion test

Bacterial cultures were grown overnight, as described above, and diluted 500-fold in fresh medium until growth reached an OD_600nm_ of 0.4-0.8. The cultures were plated on LB-agar or BHI-agar, according to the bacterial strain. Lyophilized uperin 3.5 peptide powder was dissolved in ultra-pure water to a final concentration of 50 mg/ml. The concentrated peptide solution was loaded on blank antimicrobial susceptibility discs (Oxoid, UK). Discs loaded with ultra-pure water served as controls. The discs were gently placed on bacteria-plated agar and incubated overnight. Bacterial growth on the agar plate results in an opaque surface, which is inhibited by the antimicrobial substance radially diffusing from the disc. Thus, the radius of the clear agar around the disc enables evaluation of the antibacterial propensity of the tested drug (uperin 3.5).

### Determination of the minimal inhibitory concentration (MIC)

*M. luteus* cultures were grown overnight, as described above, and diluted 500-fold in fresh medium, until growth reached an OD_600mn_ of 0.4-0.8. Uperin 3.5 was dissolved to a concentration of 10 mM in ultra-pure water and diluted to 0.5 mM in growth media (LB). Two-fold serial dilutions of uperin 3.5 in LB were prepared in a sterile 96-well plate, to obtain concentrations ranging from 1 μM to 60 or 125 μM. Wells containing growth medium and bacteria served as positive controls and wells containing growth media only served as negative control (blank). Bacterial growth was determined by measuring OD_600_ after a 24 h incubation at 30 °C, with 250 rpm shaking. The experiment was performed in triplicates or more. All triplicate values were averaged, and appropriate blanks were subtracted. The entire experiment was repeated at least three times on different days. The MIC of each sample was determined as the lowest concentration of peptide in which no growth was observed after a 24 h incubation. The MIC of the heated fresh sample was determined as described above, but after a 10 min, 60 °C heat-shock of uperin 3.5 freshly dissolved solution. The MIC of the incubated samples were determined as described, by incubating uperin 3.5 solution for 3 days, and a 10 min, 60 °C heat-shock. In all cases of heat-shock, samples were allowed to cool to RT before the assay was performed.

### Scanning electron microscopy (SEM)

*M. luteus* was grown overnight as described above and diluted 50-fold in fresh medium, until growth reached an OD_600nm_ of 0.4-0.8. The culture was centrifuged at 4000 rpm and bacterial cells were washed three times with 10 mM potassium phosphate buffer, containing 150 mM NaCl, pH 7.3 (herein, buffer). The pellet was re-suspended in the buffer and diluted to an OD_600_ of 0.2. Pretreated uperin 3.5 was dissolved to a concentration of 10 mM in ultra-pure water and immediately diluted to 1 mM in the buffer. The peptide was mixed with the bacterial suspension, to a final concentration of 2 μM or 6 μM. Samples were incubated at 30 °C, without shaking, for 24 h. Incubated bacteria (5 μl) were dispersed on 5×5 mm silicon wafers (Ted Pella, USA) that had been prewashed with ethanol, glow-discharged (PELCO easiGlow, Ted Pella) and pretreated with poly-L-lysine 0.1 % solution (30 μl, RT, 20 min). After allowing the samples to air dry for a few minutes, they were scanned with a Zeiss ULTRA plus field emission gun scanning electron microscope, operated at 1.3 kV (up to 80x magnification) using either the InLens or SE2 detectors.

### Fluorescence microscopy

Pre-treated FITC-labelled uperin 3.5 was dissolved to a concentration of 10 mM in ultra-pure water and immediately diluted to 1 mM in 10 mM potassium phosphate buffer, containing 150 mM NaCl, pH 7.3. Rhodamine-labeled DOPE:DOPG SUVs were prepared as described above and mixed with FITC-labeled uperin 3.5 solution to a final concentration of 10 μM in a 1:3 peptide: lipid molar-ratio. *M. luteus* was grown overnight, as described above, and diluted 50-fold in fresh medium, until growth reached an OD_600nm_ of 0.4-0.8. The culture was centrifuged at 4000 rpm and pellets were washed three times in buffer. The pellet was re-suspended in buffer and diluted to an OD_600_ of 0.1. FITC-labeled uperin 3.5 solution was mixed with *M. luteus* to a final concentration of 10 μM. Propidium iodide (PI) solution was added to a final concentration of 0.1 mg/ml. A sample of *M. luteus* and a sample of FITC-labelled uperin 3.5 (10 μM), each mixed with propidium iodide served as controls. A droplet (5 μl) of each sample were applied to a glass microscope slide and covered with a coverslip, and then examined under an inverted fluorescence microscope (Inverted Leica DMI8), using both transmitted light, a green filter with 460-500 nm excitation and 512-542 nm emission wavelengths to view FITC-labelled uperin 3.5, and a Texas Red filter with 540-580 nm excitation and 592-668 nm emission wavelengths, to view the rhodamine-labelled SUVs and PI. Images of fluorescence and bright-field measurements were captured and edited via the Leica DFC7000 Camera and Application Suite X (LAS X) and rendered using ImageJ.

## Acknowledgments

We wish to thank Sunny Singh for initiating fibrillation experiments and crystallization. We acknowledge guidance and technical support provided by Yael Pazy-Benhar and Dikla Hiya at the Technion Center for Structural Biology (TCSB). We acknowledge support from Yaron Kauffmann from the MIKA electron microscopy center of the Department of Material Science & Engineering at the Technion, Na’ama Koifman from the Russell Berrie Electron Microscopy Center of Soft Matter at the Technion, Nitsan Dahan and Yael Lupo-Haber from the Life Science and Engineering Infrastructure Center, Rachel Edrey from the Chemical and Surface Analysis Laboratory, at the Department of Chemistry, all at the Technion, Israel. This research was supported by Israel Science Foundation (grant no. 560/16), Israel Ministry of Science, Technology & Space (grant no. 78567), U.S.-Israel Binational Science Foundation (BSF) (grant no. 2017280), BioStruct-X, funded by FP7, and the iNEXT consortium of Instruct-ERIC. The synchrotron MX data collection experiments were performed at beamlines ID29, ID23-EH2 and ID30A-3 / MASSIF-3 at the European Synchrotron Radiation Facility (ESRF), Grenoble, France, and at beamline P14, operated by EMBL Hamburg at the PETRA III storage ring (DESY, Hamburg, Germany). We are grateful to the teams at ESRF and EMBL Hamburg.

---

**Figure S1.**
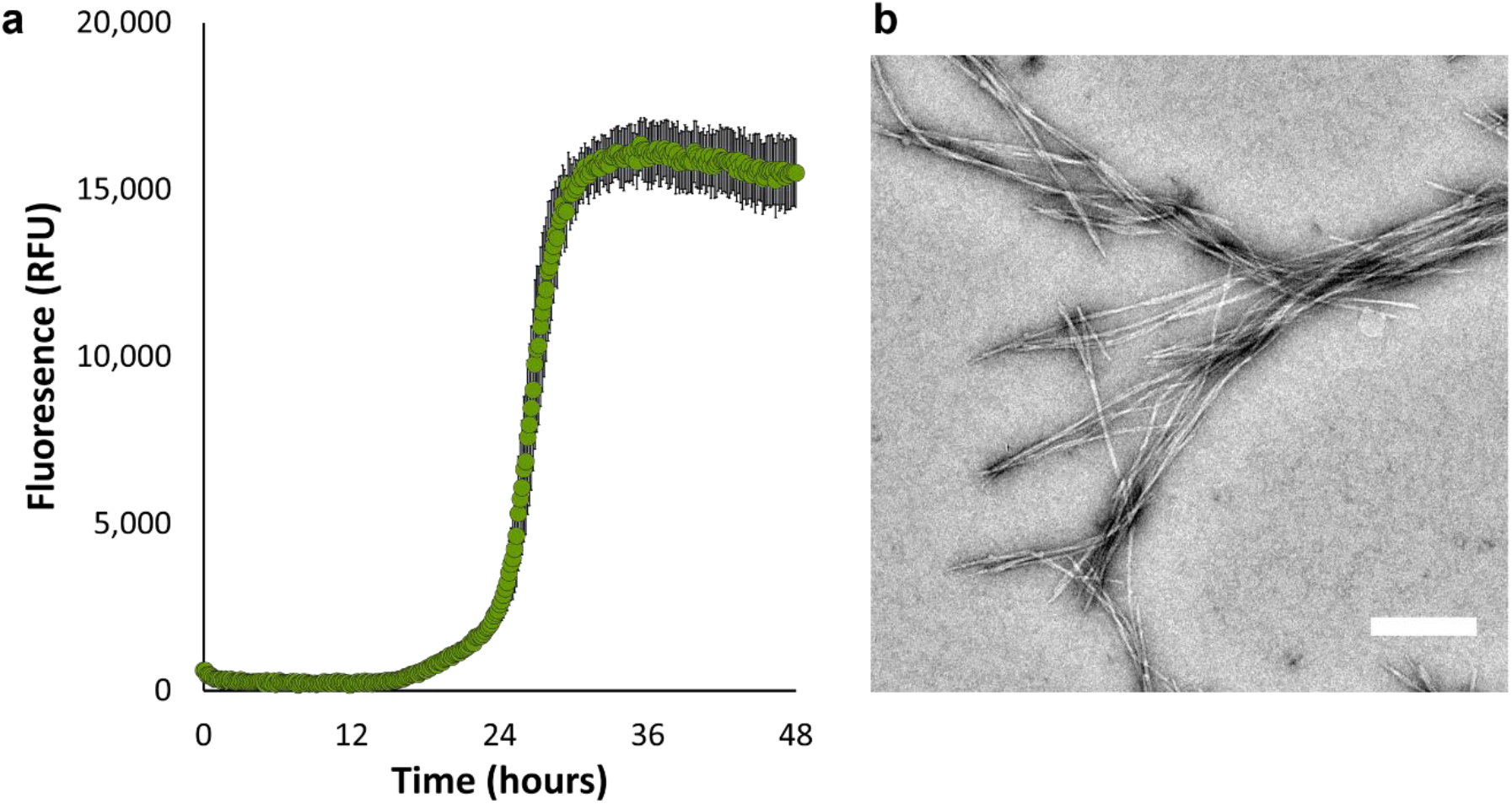
Uperin 3.5 self-assembles to form amyloid fibrils that bind thioflavin-T. (**a**) Fibrillation kinetics of 100 μM uperin 3.5 monitored by thioflavin-T (ThT) binding. Uperin 3.5 showed rapid fibrillation, following a lag time of about 16 h. The graph shows mean fluorescence readings of triplicate ThT measurements. Error bars represent standard errors of the means. (**b**) A negatively stained transmission electron micrograph demonstrating elongated fibrils of 100 μM uperin 3.5 incubated for 6 days. Scale bar represents 300 nm.

**Figure S2.**
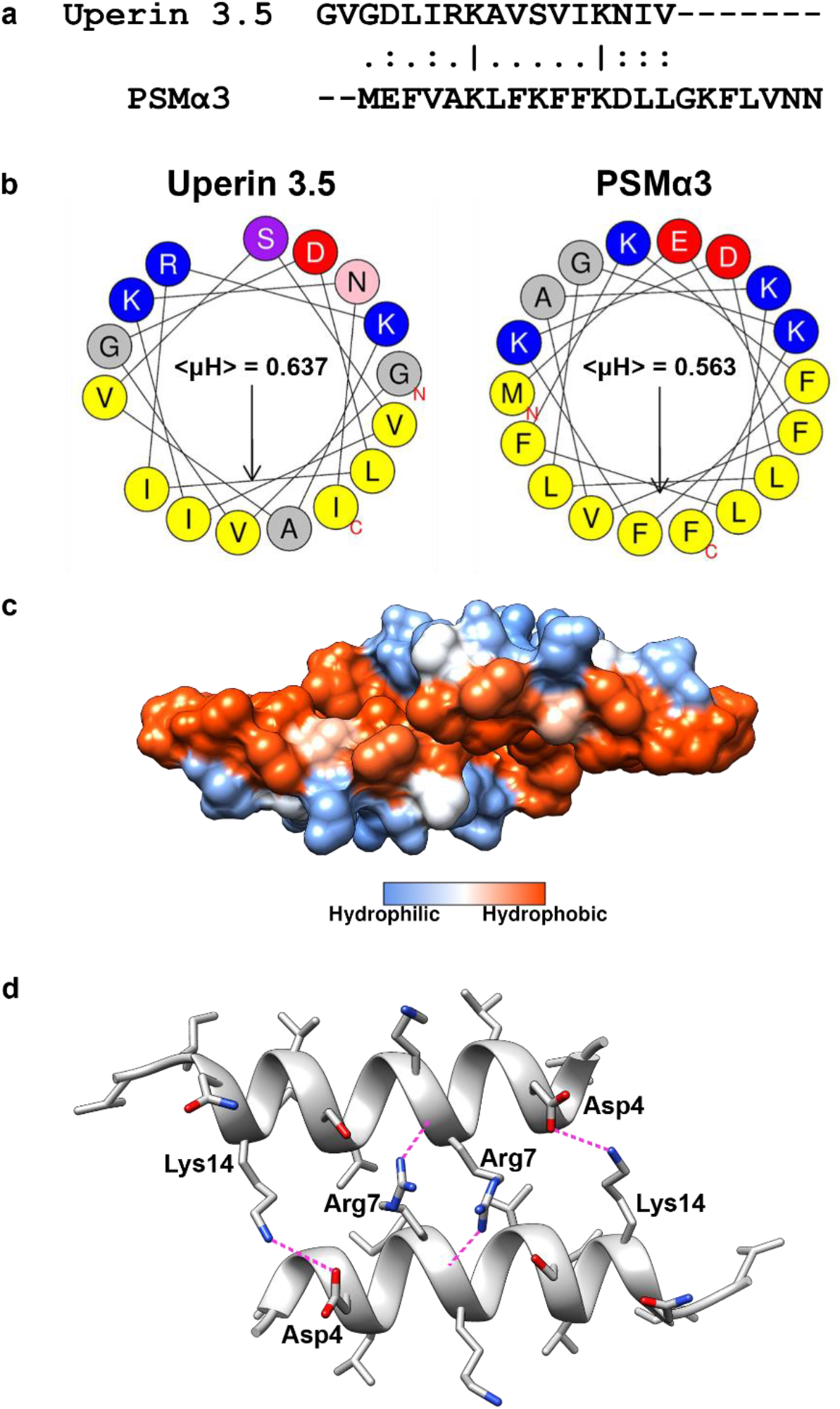
Uperin 3.5 forms cross-α amyloid fibrils stabilized by hydrogen bonds. (**a**) Uperin 3.5 and PSMα3 global sequence alignment generated using EMBOSS^1^, wherein a dot indicates low or no similarity between the residues, a colon indicates physicochemically similar residues, and a vertical line indicates identical residues. (**b**) Helical wheels of uperin 3.5 and PSMα3 generated using Heliquest^2^. The N- and C-terminal residues are marked in small red letters, the arrow represents the hydrophobic moment <μH> and its value is written inside the wheel, hydrophobic residues are colored yellow, negatively charged residues are colored red, positively charged residues are colored blue, glycines and alanines are colored grey, serine is colored purple and asparagine is colored pink. (**c**) Uperin 3.5 forms a hydrophobic interface between mating sheets forming the fibril. The view is down the fibril axis, showing six layers of peptides, shown in surface representation, and colored by hydrophobicity according to the scale bar. (**d**) Polar, inter-helical interactions along the uperin 3.5 fibril. The view is perpendicular to the fibril axis, showing only two helices (in grey) along one sheet. The peptide backbone is shown in ribbon, side chains are shown as sticks and heteroatoms are colored according to atom type (nitrogen in blue and oxygen in red). Polar interactions are depicted with dashed lines colored pink and the interacting residues are labeled.

**Figure S3.**
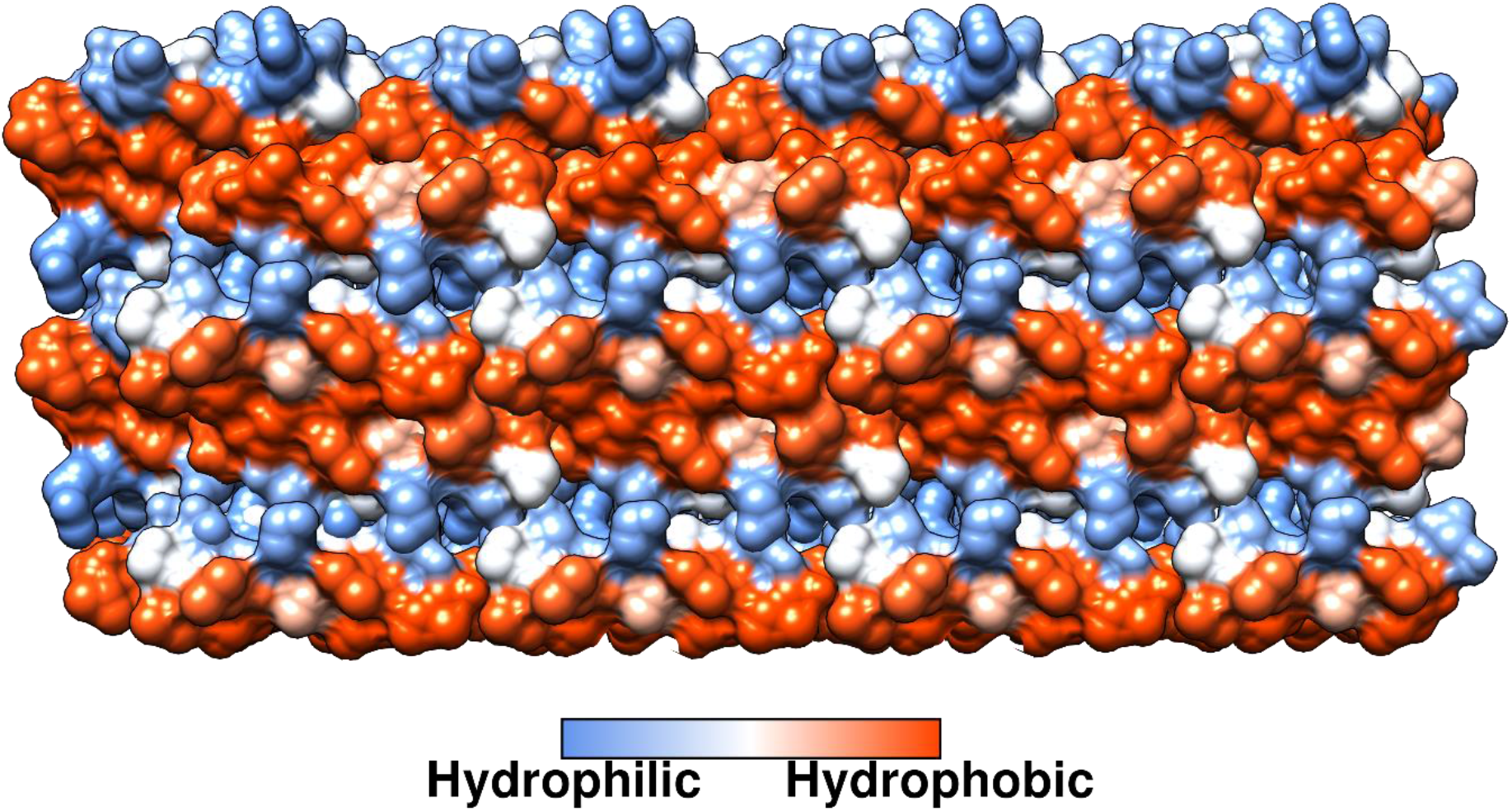
Uperin 3.5 high-order crystal packing showing alternating hydrophilic and hydrophobic interfaces. The high-order crystal packing of uperin 3.5 is depicted down the fibril axis, with continuous rows of amphipathic α-helices shown tightly packed via their hydrophobic faces into mating sheets, forming alternating hydrophilic and hydrophobic interfaces. Peptides are shown in surface representation, and colored by hydrophobicity, according to the scale bar.

**Figure S4.**
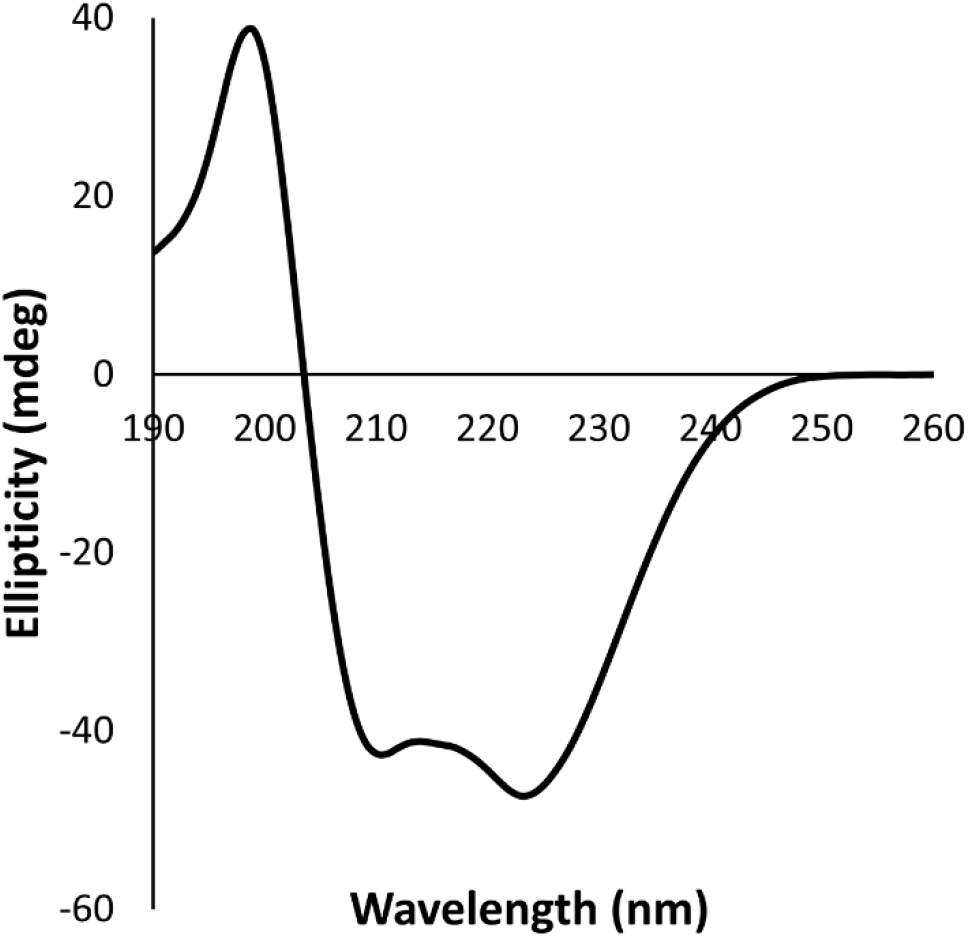
Secondary structure of PSMα3 cross-α fibrils in solid-state CD. Solid-state CD (ssCD) spectrum of PSMα3 thin films, indicating the presence of cross-α fibrils. The ssCD spectrum showed a deeper minimum at 222 nm as compared to the 208 nm minimum, suggestive of interacting α-helices, further supporting the proposed cross-α architecture of the fibrils^3,4^. This resembled the α-helical ssCD spectrum of uperin 3.5 fibrils pre-formed in the presence of 1,2-dioleoyl-sn-glycero-3-phosphoethanolamine and 1,2-dioleoyl-sn-glycero-3-phospho-(1’-rac-glycerol) (DOPE and DOPG) SUVs (Figure 2B).

**Figure S5.**
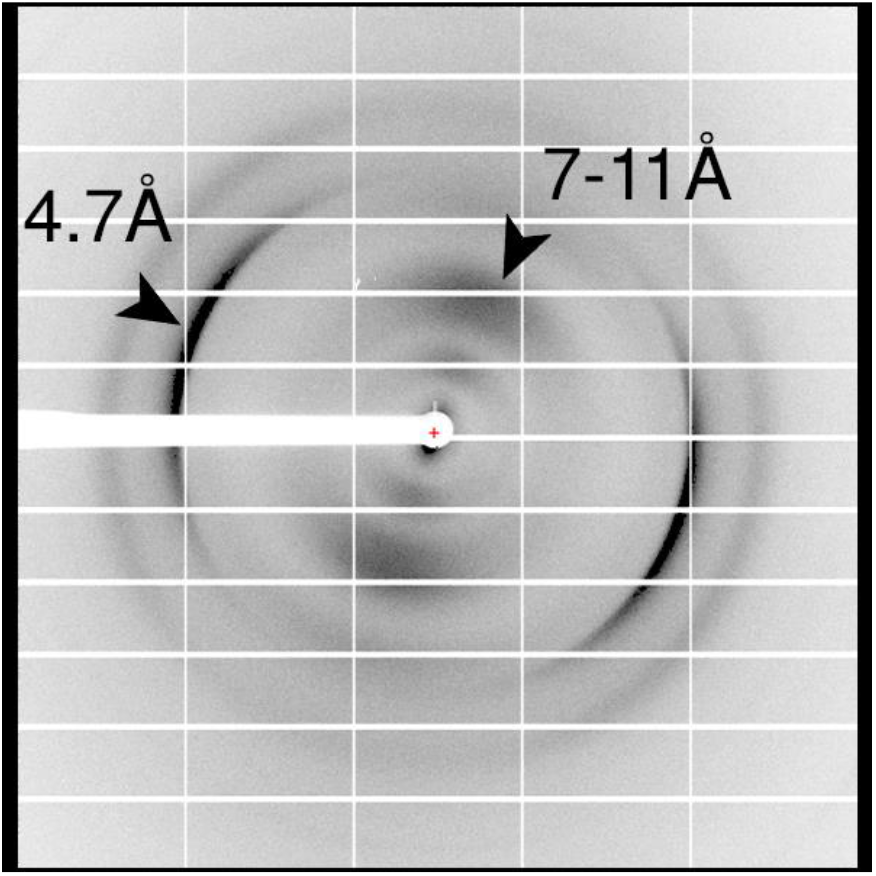
X-ray fiber diffraction of uperin 3.5. X-ray fiber diffraction of uperin 3.5 showed a typical cross-β signature^5^, with major diffraction arches at 4.7 Å (sharp), and orthogonal reflection at 7-11 Å (diffused).

**Figure S6.**
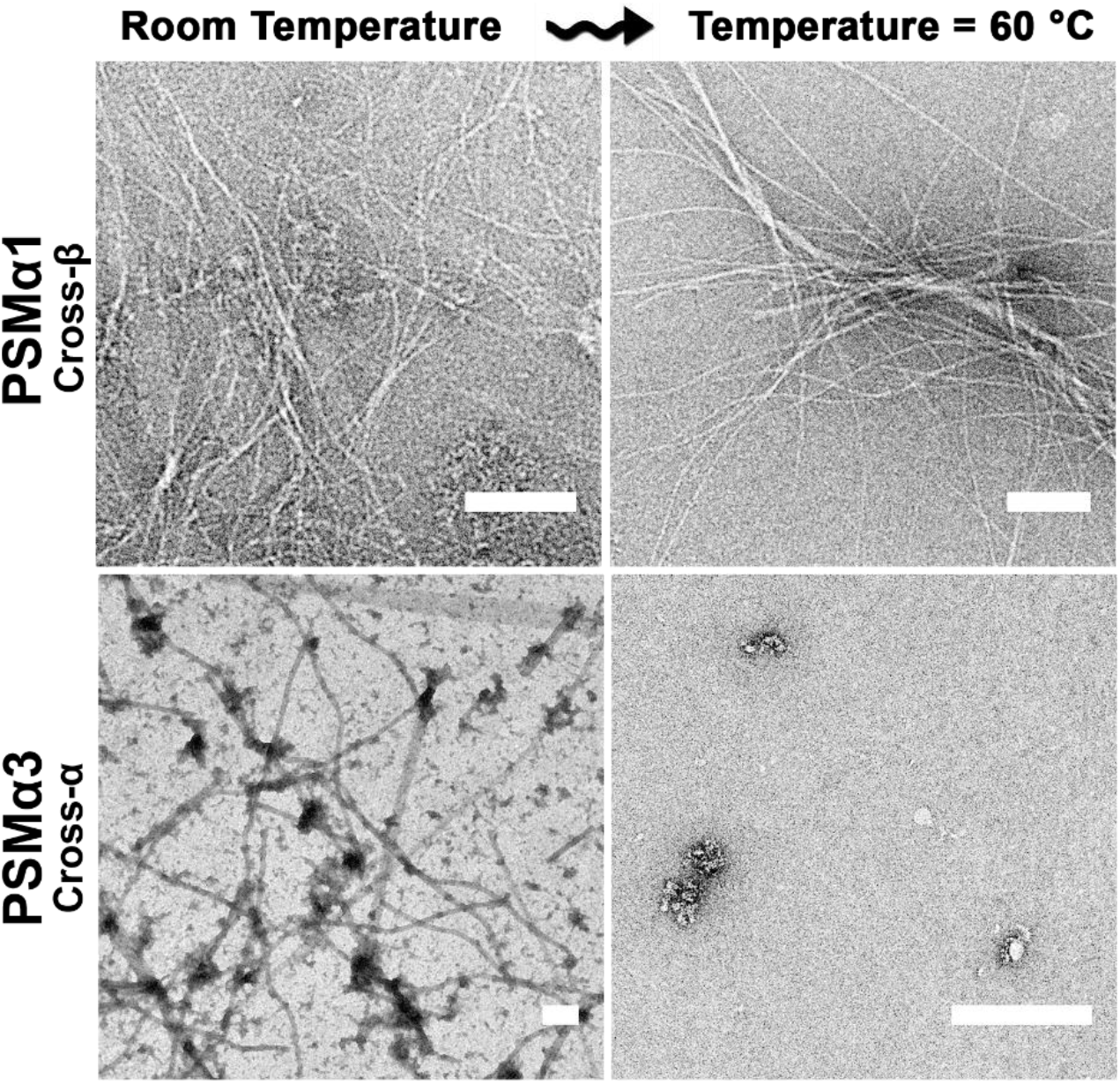
Thermal stability of amyloid fibrils with different architectures. The cross-β-forming PSMα1 showed more thermostable fibrils, in contrast to the cross-α-forming PSMα3 fibrils, which dissolved upon heating to 60°C. The thermostability of uperin 3.5 chameleon cross-α/cross-β fibrils is shown in figure 3B. Samples at 10mM were incubated for 2 days at room-temperature, diluted 10-fold, and fixed on electron microscopy grids, and fixed again immediately after a 10-min, 60°C heat shock treatment. For PSMα1, scale bars represent 200 nm; for PSMα3, scale bars represent 500 nm.

**Figure S7.**
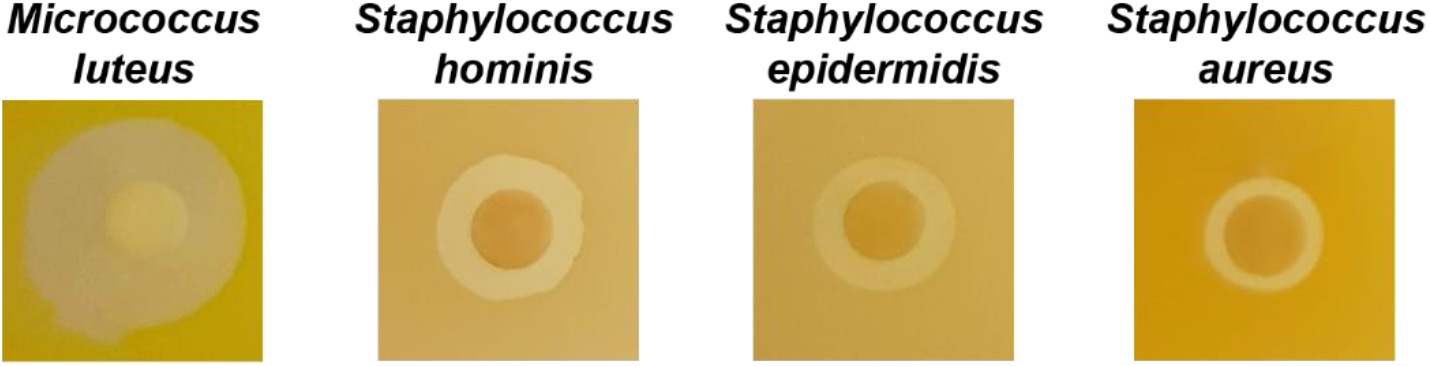
Uperin antibacterial activity against different Gram-positive pathogens. Antibacterial activity of uperin 3.5 against different Gram-positive pathogens tested via a disc (agar) diffusion assay. In this assay, the test agent diffuses into the agar and growth inhibition of the tested microorganism is evaluated. Of note, the different bacterial media used result in different agar color. Uperin 3.5 potently affected *M. luteus* growth in comparison to other strains tested.

**Figure S8.**
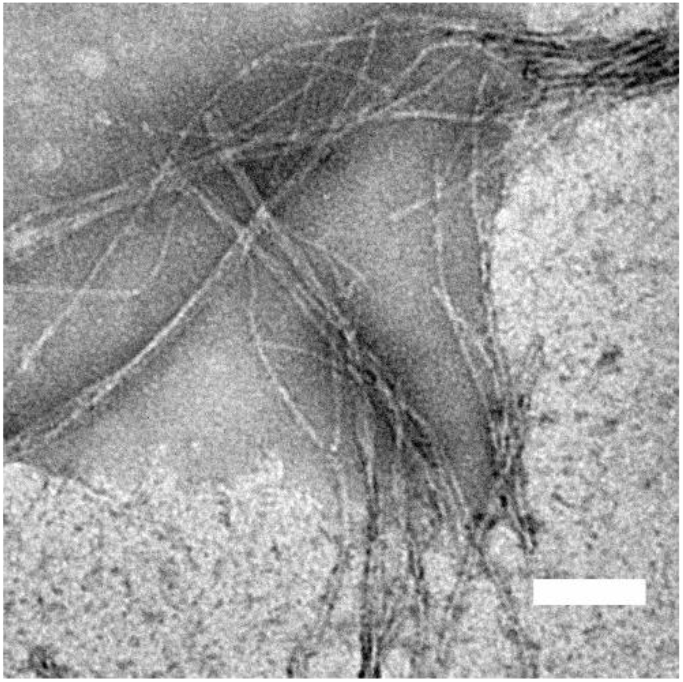
FITC-labelled uperin 3.5 forms fibrils. Electron micrograph demonstrating negatively stained, elongated fibrils of FITC-labelled uperin 3.5 incubated at 1 mM for 5 days. Scale bar represent 100 nm. Unlabeled uperin 3.5 fibrils formed following a 2-day incubation at the same concentration, are shown in Figure 3.

**Figure S9.**
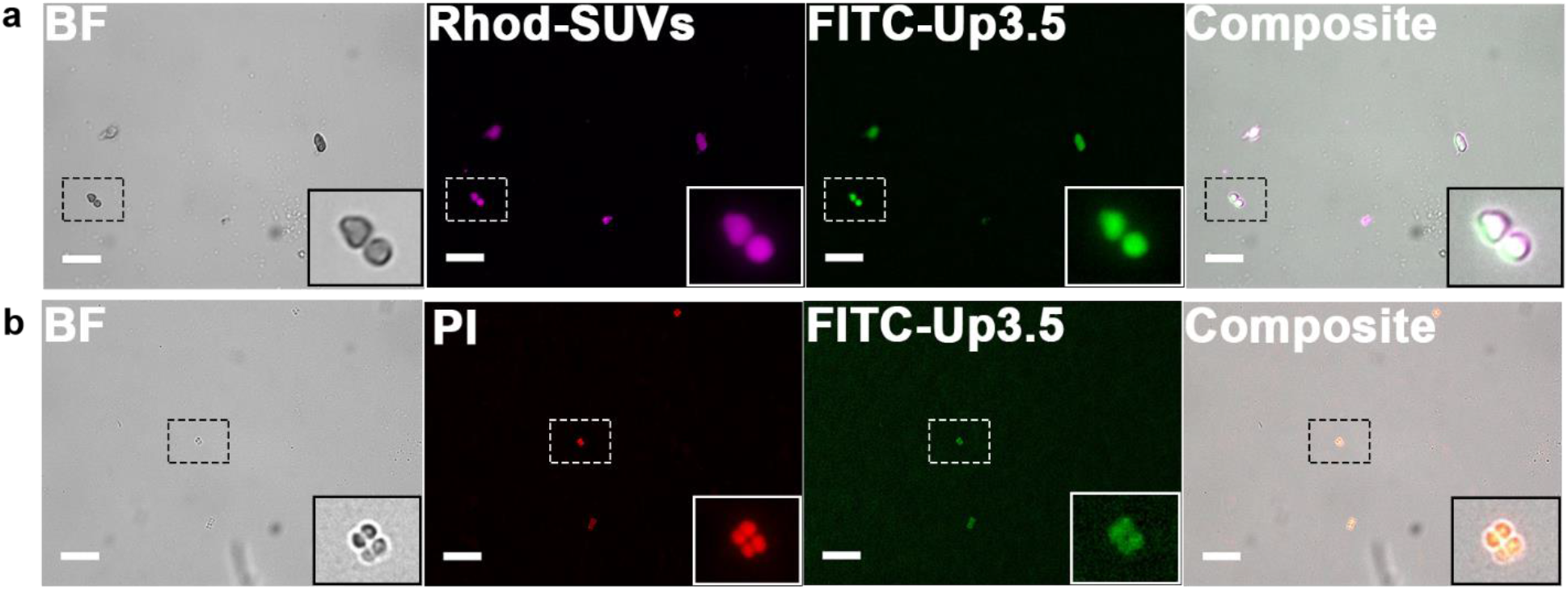
Fluorescence microscopy images of uperin 3.5 foci co-localizing with SUVs or bacterial cells and inducing cell death. Fluorescence microscopy images of 10 μM FITC-labeled uperin 3.5 mixed with rhodamine-labelled DOPE:DOPG SUVs (**a**) or *M. luteus* cells (**b**). In each image, the area marked by a dashed rectangle is enlarged in the insert in the bottom right corner. In the composite images (right panels), the bright field (BF) and FITC (green) channels were merged with either rhodamine (Rhod, purple) (a) or propidium iodide (PI; red) (b). FITC-labelled uperin 3.5 accumulated in or inside the SUVs (a), or *M. luteus* cells, inducing cell death, as shown by the uptake of propidium iodide (PI) (b). Scale bars represent 20 μm for all images.

**Table S1.**
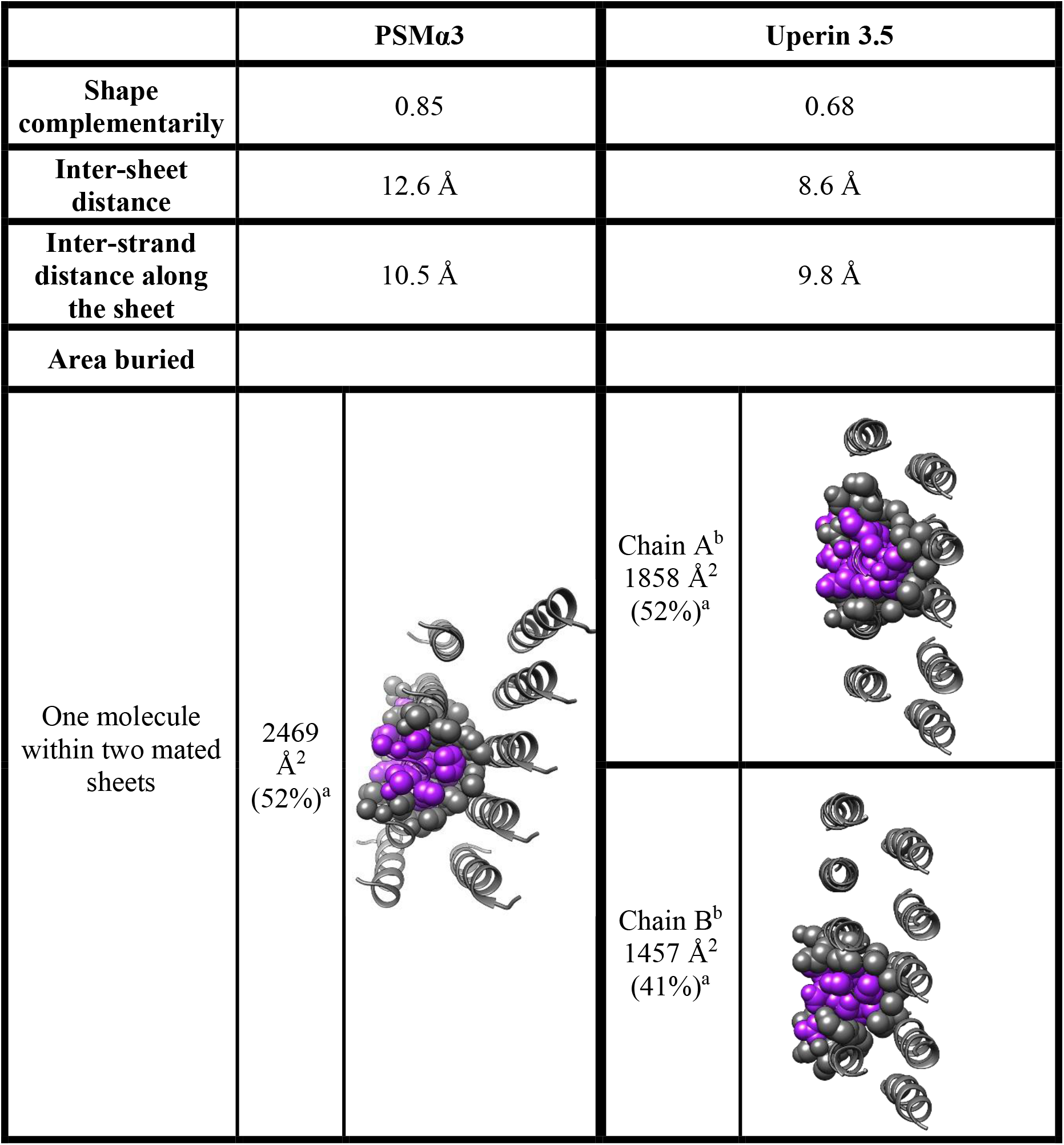
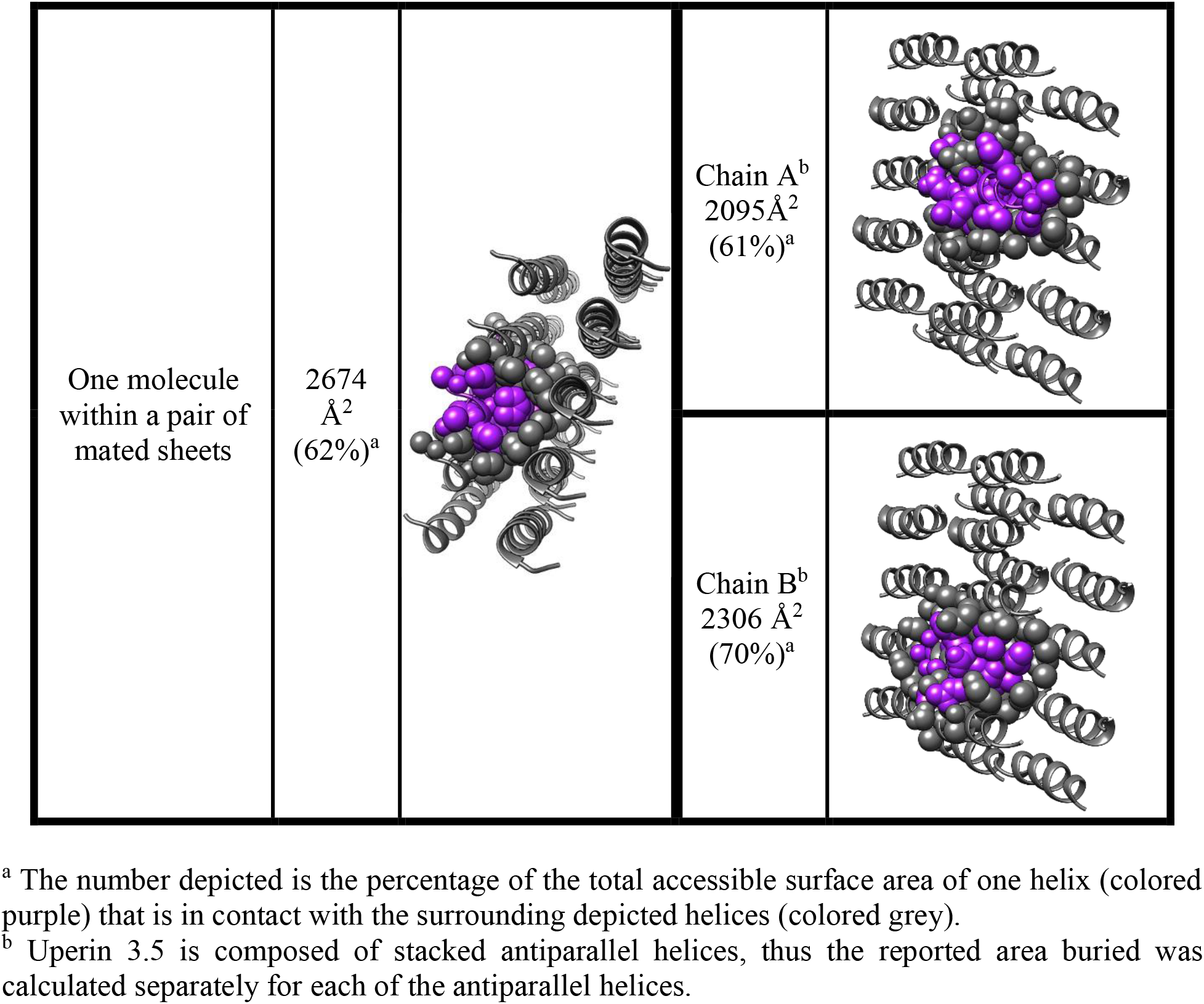
Features of the uperin 3.5 cross-α structure as compared to PSMα3.

| | PSMα3 | | Uperin 3.5 | |
| --- | --- | --- | --- | --- |
| **Shape complementarity** | 0.85 |  | 0.68 |  |
| **Inter-sheet distance** | 12.6 Å |  | 8.6 Å |  |
| **Inter-strand distance along the sheet** | 10.5 Å |  | 9.8 Å |  |
| **Area buried** |  |  |  |  |
| One molecule within two mated sheets | 2469 Å<sup>2</sup><br>(52%)<sup>a</sup> |  | Chain A<sup>b</sup><br>1858 Å<sup>2</sup><br>(52%)<sup>a</sup> |  |
|  |  |  | Chain B<sup>b</sup><br>1457 Å<sup>2</sup><br>(41%)<sup>a</sup> |  |
| One molecule within a pair of mated sheets | 2674 Å<sup>2</sup> (62%)<sup>a</sup> |  | Chain A<sup>b</sup><br>2095 Å<sup>2</sup> (61%)<sup>a</sup> |  |
|  |  |  | Chain B<sup>b</sup><br>2306 Å<sup>2</sup> (70%)<sup>a</sup> |  |

<sup>a</sup> The number depicted is the percentage of the total accessible surface area of one helix (colored purple) that is in contact with the surrounding depicted helices (colored grey).
<sup>b</sup> Uperin 3.5 is composed of stacked antiparallel helices, thus the reported area buried was calculated separately for each of the antiparallel helices.

